# Pupil Dynamics Reflect Uncertainty-Driven Adjustments of Probability Learning

**DOI:** 10.64898/2026.02.26.708148

**Authors:** A. Greenhouse-Tucknott, C. Foucault, A. Buot, F. Meyniel

**Affiliations:** Cognitive Neuroimaging Unit, INSERM, CEA, CNRS, Université Paris-Saclay, NeuroSpin center, 91191 Gif/Yvette, France; Institut de neuromodulation, GHU Paris, psychiatrie et neurosciences, centre hospitalier Sainte-Anne, pôle hospitalo-universitaire 15, Université Paris Cité, Paris, France; Department of Experimental Psychology, University of Oxford, Oxford, United Kingdom

## Abstract

Adaptive learning in dynamic environments depends on the brain’s ability to represent and respond to uncertainty. Changes in arousal-mediated pupil diameter provide insight into the neural basis of this process. Prior work has focused on magnitude-based inference tasks and revealed that learning is primarily influenced by the detection of change points and the corresponding modulation of arousal. In comparison, the role of arousal systems in learning contexts where changes are harder to detect, such as in probability learning, remains poorly understood. Here, we combined pupillometry with a probability learning task in which human participants estimated a hidden generative probability of stimuli that changed abruptly and unpredictably. A Bayesian ideal observer model was used to estimate trial-by-trial learning rates and two uncertainty-associated factors: change point probability (the probability that the generative process has abruptly changed) and prior uncertainty (about the current beliefs on this generative process). In line with normative theory, participants relied on prior uncertainty to adjust learning rates. Pupil analyses revealed temporally distinct representations of uncertainty, with phasic dilations tracking change point probability and tonic pupil dilations tracking prior uncertainty. Mediation analyses further indicated that both phasic and tonic pupil signals partially carried the effects of change point probability and prior uncertainty on learning, respectively. Critically, high tonic arousal weakened the coupling between phasic pupil responses and change point probability, accompanied by a reduced influence of change point probability on learning. Together, our results demonstrate that probability learning in a dynamic environment is underpinned by computationally rational integration of latent uncertainty factors and its contribution to belief updating, regulated through arousal-associated responses.

**AUTHOR SUMMARY:** Probabilistic events pervade our environments (*e.g.* think about the probability you will have to wait a long time for a table at your favourite restaurant). We learn about these probabilities by integrating new observations into existing beliefs. However, probabilistic events naturally fluctuate due to chance (*i.e.* they are stochastic), creating uncertainty about the true probability. Because any new observation is only weakly informative, learning is often slow and it can be difficult to distinguish other sources of uncertainty, such as profound environmental changes (*i.e.* volatility). Yet here, we show that human learners are capable of adapting how they learn about probabilities within dynamic environments, indicating mechanisms that track different sources of uncertainty. Using pupillometry, we show that arousal systems provide one such mechanism, with distinct patterns of arousal tracking uncertainty about current beliefs and the likelihood of environmental change. Importantly, these signals interact: as overall arousal increases, uncertainty about current beliefs exerts a stronger influence on learning, while arousal responses associated with detecting environmental change become less linked to learning. These findings suggest that arousal systems dynamically regulate which sources of uncertainty are prioritised when updating probability beliefs in changing environments.

## INTRODUCTION

The capacity to learn and predict future outcomes from past experiences is of paramount importance for adaptive behaviour [1]. Learning depends critically on how sources of uncertainty are represented and managed [2]. Previous studies have shown that humans do regulate how much they learn [3–8], with this adaptation depending on the level of stochasticity (inherent variability of outcomes) and volatility (changes in the environment), two environmental factors that cause uncertainty [9,10]. To illustrate, waiting times at a busy restaurant may fluctuate because of many factors. Experience allows one to learn the typical range of waiting times, which requires all the more observations than variability is higher. In contrast, an unusually long wait may indicate a fundamental change such as new management or staffing, which requires a prompt learning reset.

The rate of learning controls the extent to which beliefs are updated in response to new observations, and it is adapted to different types of uncertainties. Lower learning rates give greater weight to past experience, favouring stability of estimates, whereas higher learning rates give greater weight to recent observations, favouring learning reset. Learning rates can be studied as explicit parameters of computational models [11,12]. Alternatively, an apparent learning rate can be measured as the ratio between the belief update and the prediction error [5,13]. We use “apparent” here because this metric is agnostic of the actual learning algorithm of the agent. To understand how learning rates are adjusted trial-by-trial, normative (Bayesian) models have been adopted to estimate latent uncertainty variables and impact on learning. These studies indicate that learning is shaped by at least two forms of uncertainty: prior uncertainty, reflecting imprecision in existing beliefs, and change-point probability, reflecting the likelihood that an abrupt environmental change has occurred [14–20].

The neural mechanisms governing the uncertainty-dependent regulation of learning remain largely unknown. Neuromodulatory systems have long attracted attention as putative mechanisms [9,21–23]. Used as a non-invasive proxy of central arousal state [24–28], in humans there has been a heavy leverage of nonluminance-dependent, pupil dilations to understand how uncertainty may be represented by neuromodulatory systems to shape cognitive processes such as learning and decision-making [18,29–31]. During learning, transient, phasic pupil dilations to new observations consistently scale with computational markers of surprise and environmental change [4,32,30,33,34,18]. Indeed, Nassar and colleagues [18] demonstrated that these phasic pupil responses track change point probability during a magnitude inference task. The authors further demonstrated a causal link between these phasic pupil responses and learning rate by experimentally increasing arousal, which enlarged pupil diameter and enhanced learning. Notably, this evoked effect depended on baseline pupil diameter. Changes in baseline (*i.e.* tonic) pupil size may reflect another form of uncertainty-weighting, as increasing evidence has linked baseline pupil changes with tracking prior belief uncertainty [18,34–38]. Phasic- and tonic-associated pupil effects can be experimentally dissociated [39], offering the possibility that distinct neural structures are engaged for learning regulation which are sensitive to the prominence of environmental uncertainty parameters.

Our current understanding of how latent uncertainty factors drive trial-by-trial changes in apparent learning rates is almost exclusively informed by magnitude learning tasks. These tasks have utilised magnitude-based quantities, such as spatial positions [17,19,20] or numerical values [18]. In such cases, apparent learning rates are highly determined by dynamics in change point probability [14] which, as introduced, are associated with phasic activations of arousal [18]. However, magnitude learning is fundamentally different to probability learning (i.e. learning the probability of discrete outcomes). Specifically, each new observation during probability learning carries much less information about the inferred quantity compared to magnitude learning, meaning that it is much harder to detect change points and appropriately update beliefs. This difference promotes a critical shift in how apparent learning rates are driven: probability learning should place less emphasis on the change point probability associated with a new observation and instead rely more on prior uncertainty [14].

It remains unknown whether the representations of latent uncertainty factors across tonic and phasic arousal states support the trial-by-trial regulation of apparent learning rates during probability learning. To address this outstanding question, we employed a previously developed probability learning task [14] to continuously capture participants’ apparent learning rate, coupled with pupilometry to concomitantly assess changes in pupil diameter. The Bayesian optimal solution to the inference problem [34] was applied to not only provide a reference comparison for the analysis of participants’ behavioural estimates, but also extract trial-by-trial latent uncertainty factors. These uncertainty factors were then used to evaluate drivers of learning regulation and the modulation of pupil dilatory responses. Overall, our results support the hypothesis that during probability learning, prior uncertainty dominates in the regulation of learning. Prior uncertainty and change point probability are separately encoded within tonic and phasic pupil signals, respectively, and these identified pupil signals, at least in part, mediate the effect of the associated latent uncertainty factor in the regulation of learning, providing a physiological mechanism supporting belief updating during probability inference.

## RESULTS

The probability learning task required participants to continuously infer the value of a hidden state - in this instance, the generative stimulus probability - from a sequence of stimuli [14]. We used a cover story for the task to improve comprehension. The cover story described a spinning roulette wheel which was divided into two colours (*i.e.* blue and yellow). The proportional distribution of the colour blue corresponded to the hidden generative probability. Participants were tasked with trying to estimate the proportion of blue (*vs.* yellow) on the wheel as accurately as possible based on a sequence of the two colours presented to them. The sequence presented was described as the result of repeatedly spinning the wheel. Participants continuously updated their estimate of the hidden proportion after each new observation by moving the marker on a semi-circular slide tracker (*Fig 1*). Each sequence consisted of 75 observations, with observations presented as a coloured circle with a white target overlaid on top. Participants were instructed to maintain a fixed gaze on the target for the accurate recording of pupil diameter to each new observation throughout the task. The semi-circular slide tracker made fixation easier (see *Methods* for more details).

**Fig 1.**
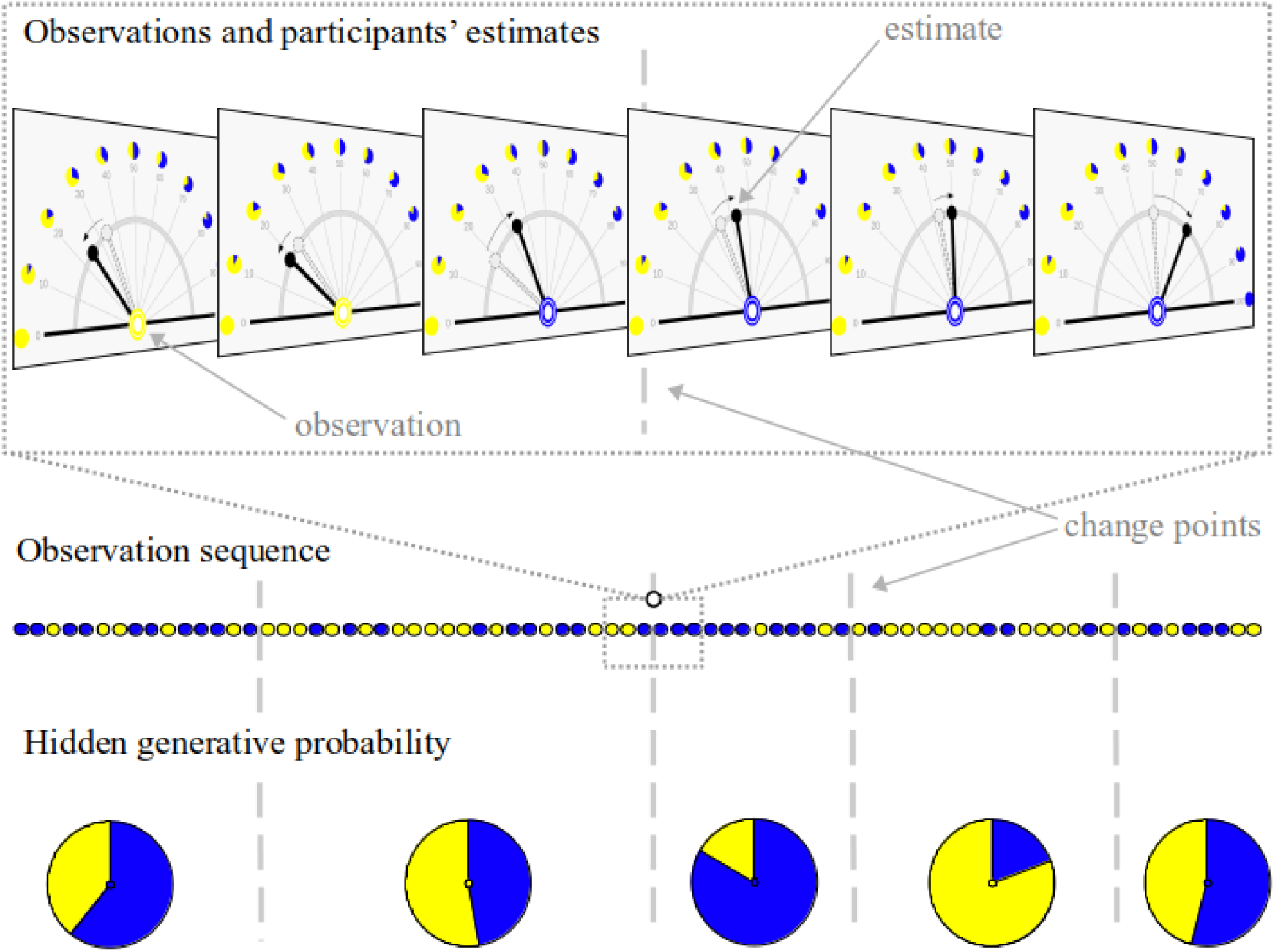
The dynamic probability learning task. Participants were shown a sequence of 75 blue and yellow balls at the fixation dot (center of the screen). They were required to continuously estimate the hidden generative probability of the colour blue (vs. yellow). To make the task more intuitive, participants were told that this probability corresponded to the proportion of blue (vs. yellow) on a fictitious spinning roulette wheel. During the sequence the hidden generative probability could change at undeclared points, referred to as change points (depicted by the dotted lines and changing proportions on the hidden generative probabilities). Change points within a sequence were sampled with constant probability and the stimulus generative probability was sampled within the range of 0.1 to 0.9. Participants reported their estimated probability by moving a marker along a semi-circle. Ticks marked every 10% of the probability with the corresponding roulette wheel depicted. This display maintains a constant distance between the cursor and the fixation dot to prevent saccades and in order to record pupil size throughout. The zoom depicts the behavior of a mock participant across 6 trials containing a change point half through. Participants completed 22 sequences (task sessions). The colours presented in this figure do not accurately reflect the colours presented in the task and are displayed as such simply to differentiate and emphasise the two different outcomes.

During each sequence, the generative probability (and thus the proportion of the two colours on the fictitious roulette wheel) could change at unpredictable times, which were referred to as change points. Participants were aware of the possibility that the proportional distribution of the colours could change, but were not signalled when they happened during the observation sequence. The generative probability was uniformly sampled within the range of 0.1 and 0.9, preventing long streaks of observations where no updates are required. The magnitude of the change point was constrained, to avoid changes that are too subtle to be perceived (see *Methods* for more details). As described previously, the task is engaging and participants are capable of tracking changes in the generative probability [14].

### Participants continuously adapt estimates in a way that closely follows optimal inference

We employed a Bayesian ideal observer model to provide an optimal solution to the inference problem as a principled point of comparison [40]. The model returns a posterior distribution of the hidden generative probability that is updated upon each new observation (*Fig 2B*). We also extracted the dynamics of two latent, uncertainty-associated factors: prior uncertainty and change point probability (*Fig 2F and 2G*). Here, prior uncertainty is quantified as the standard deviation of the hidden probability estimated just before the current observation, while change point probability reflects the probability a change in the hidden value has occurred since the last observation - conceptually akin to the magnitude of prediction error (for further specification see *Methods*).

**Fig 2.**
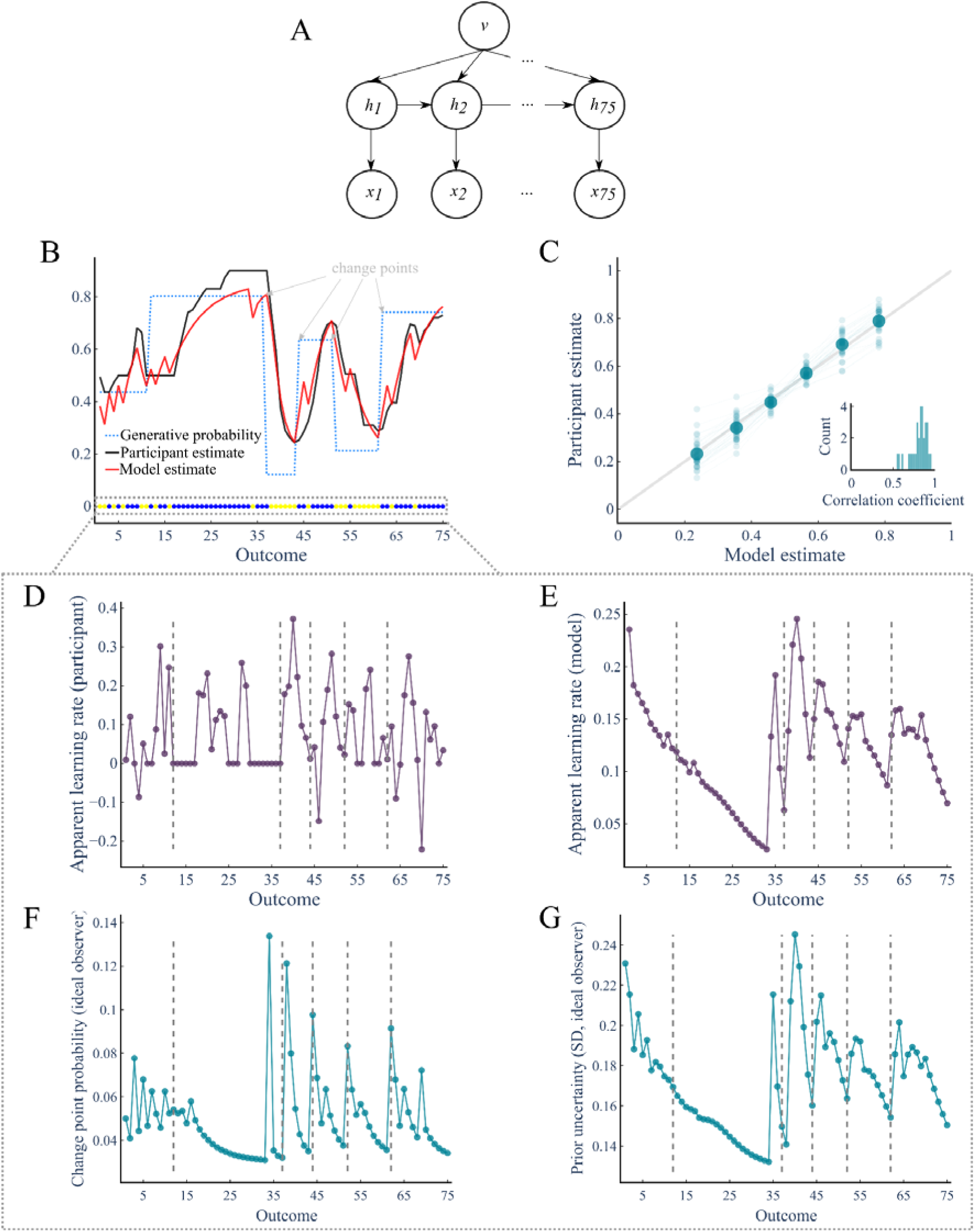
Subjective probability estimates and the ideal observer model. **(A)** A Bayesian ideal observer model was adopted to provide an optimal solution to the inferential problem. Panel A, describes the dependencies within the model. Based on a Markovian process, the model infers the posterior distribution of the (hidden) generative probability (*h_t_*) given previous observations (*x_1:t_*), assuming the true level of volatility (ν) and that the new generative probability is sampled uniformly from a proposal distribution. **(B)** An example stimulus sequence is represented by the coloured circles at the bottom of the top plot. The dotted blue line represents the (hidden) generative probability across the sequence, demonstrating a number of change points over the sequence. The black and red lines represent the estimates from the example participant and model, respectively. **(C)** Accuracy of participants’ estimates for each observation compared to model estimates (see text). For illustration purposes, the data were binned into 6 equal quantiles of model estimate and averaged within-participants. Small points and lines are the average for each participant, with the larger points reflecting the group mean. The dotted line represents the identity line. Inset displays a histogram of the subject-level coefficients (estimated without binning data). **(D and E)** Apparent learning rates for both participants and the model were derived on each observation from the degree beliefs were updated in proportion to the level of prediction error based on a simple delta rule. **(F and G)** From the model, latent uncertainty-associated factors (*i.e.* prior uncertainty and change point probability) were also extracted for each observation. Thin dotted vertical lines in panels **D-G** represent the first observation after a change point.

Evaluating participants’ performance across the entirety of the task, we found stable performance across sequences (see *Supplementary Material, Fig S1*). Our primary analysis of performance focussed on how well participants’ estimates compared to the optimal solution. Participants displayed a strong ability to accurately estimate the hidden generative probability given the relationship with the mean of the model’s posterior distribution (mean ± s.e.m; Pearson’s correlation = 0.81 ± 0.02; *t*_(30)_ = 44.72, *p* ≈ 1 × 10^-28^; *Fig 2C*; for more details on the relationship see *Supplement Material, Table S1*). Consistent with instructions, participants updated their estimate on most trials (mean ± s.e.m; 62.3 ± 3.49%). The frequency of updates was not associated with the strength of relationship between participant and model estimates (Pearson’s correlation: 0.003, p = 0.987). The participants’ apparent learning rate significantly correlated with the apparent learning rate derived from model estimates (Pearson’s Correlation = 0.07 ± 0.02; *t*_(30)_ = 4.43, *p* = 0.0001; see *Supplement Material, FigS2 and Table S1*). This correlation, although reliable across participants, was weak in magnitude; which may reflect noise in apparent learning rates at the trial level. This noise may, at least in part, be explained by trial-specific “lags” in how participants’ updates their beliefs, which we highlight in analysis presented below.

### Prior uncertainty dominates in driving apparent learning rate adjustments

The optimal solution prescribes that learning should not be constant but dynamically adjusted from trial to trial. Around change points, participants displayed a protracted increase in their apparent learning rate, which was significantly different from baseline learning rates (i.e. before the change point) from the third to the eleventh observation after a change (*Fig 3A*). This pattern conformed with normative theory (*Fig 3A insert*) (group-level correlation between participants apparent learning rate and model-derived learning rates around change points: group-level Pearson’s correlation = 0.95; participant-level Pearson’s correlations: 0.48 ± 0.06; *t*_(30)_ = 7.88, *p* ≈ 1 × 10^-9^), replicating a previou study [14].

**Fig 3.**
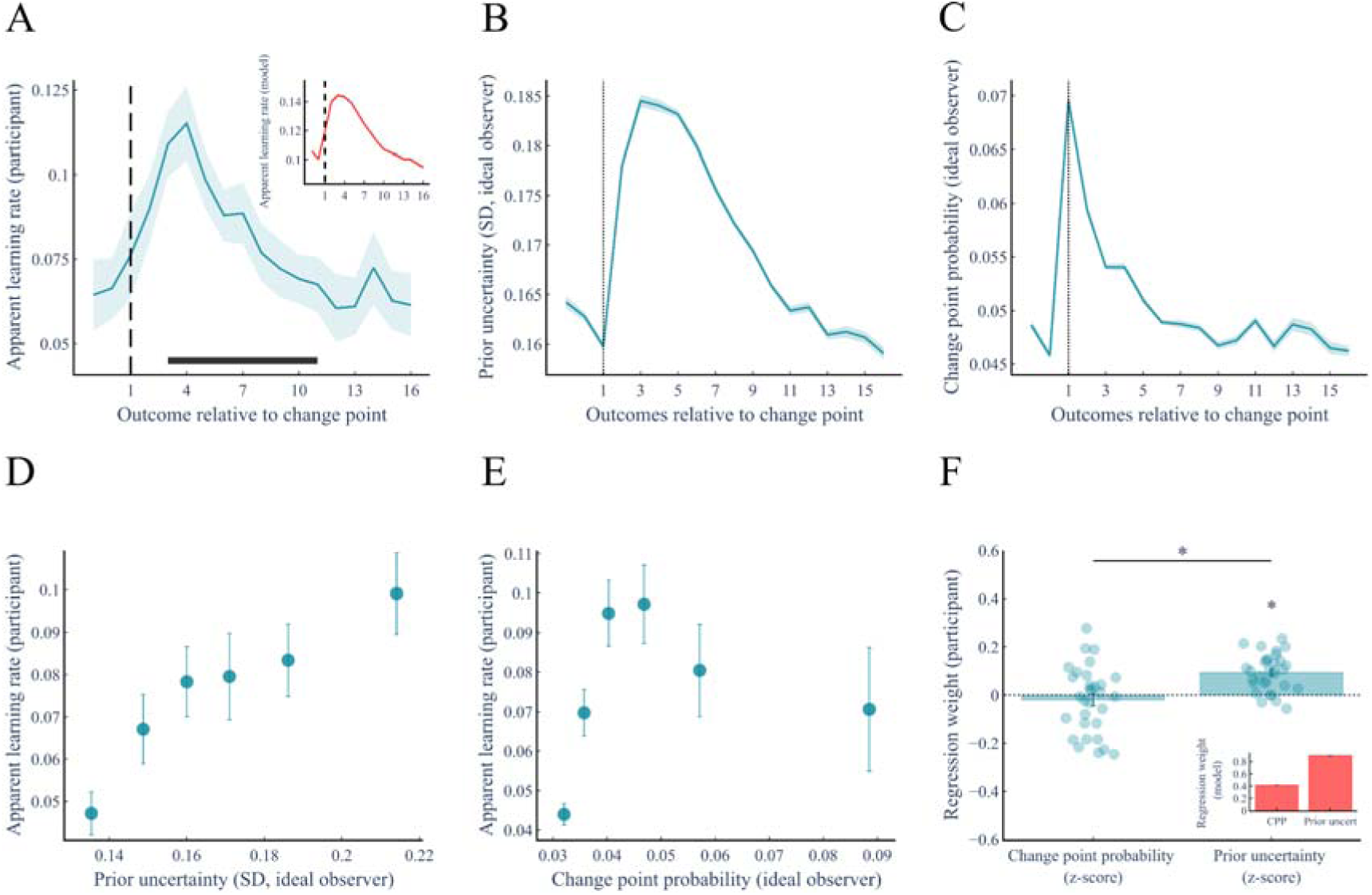
Uncertainty factors guide the dynamic adjustment of participants’ apparent learning rate. **(A)** The dynamics of participants’ apparent learning rate were examined around change points. Participants dynamically increase their apparent learning rate after a change point, with learning increased for a protracted period as new stable beliefs are formed. The horizontal line denotes the statistically significant differences compared to the pre-change point learning rate (p < 0.05, two-tailed, FWE cluster-corrected for multiple comparisons across time). The average response was similar to normative theory, shown in the inset. **(B)** Prior uncertainty parallels the change in learning rate, dramatically increasing after change points and remaining elevated for multiple trials after as new evidence is accumulated to form stable beliefs. **(C)** However, change point probability displays an increase after a change point that quickly returns back to baseline. **(D)** Across all trials, participants’ apparent learning rate increases as prior uncertainty increases (illustrated here with 6 quantiles of prior uncertainty). **(E)** For change-point probability, there was no clear linear relationship with participants’ apparent learning rate. Apparent learning rates tended to increase as change-point probability increased from low to moderate levels, but this positive association weakened and was no longer evident at higher change-point probabilities **(F)** Weights of prior uncertainty and change point probability effects on participants’ learning rates, calculated per participant by multiple linear regression. As predicted by normative theory (inset), prior uncertainty was the dominant driver of participants apparent learning rate (* represents p <0.05). The effect of prior uncertainty was greater than that observed for change point probability, which was not identified as a predictor of participant’ apparent learning rate. All plots display the mean ± s.e.m derived across all participants (n = 31).

Normative theory also prescribes that both latent uncertainty factors should play a role in determining the apparent learning rate, with prior uncertainty dominating. We initially found partial support for this prediction (*Fig 3F*). In line with normative theory, prior uncertainty was a significant driver of participants’ apparent learning rate (beta = 0.09 ± 0.01; *t*_(30)_ = 6.79, *p* ≈ 1 × 10^-7^). This could be clearly seen when examining the dynamics of prior uncertainty around change points, which paralleled the participants’ apparent learning rate (*Fig 3B*). We also observe strong graded co-variation between prior uncertainty and participants’ apparent learning rate across all trials (*Fig 3D*). The effect of prior uncertainty was significantly larger than the effect of change point probability (*t*_(30)_ = 4.48, *p* ≈ 1 × 10^-4^), as predicted by normative theory.

However, although normative theory prescribes that the effect of change point probability on the apparent learning rate should also be significant, our initial analysis did not reveal such a significant effect (beta = -0.02 ± 0.02; *t*_(30)_ = -0.80, *p* = 0.432; *Fig 3F*). The apparent learning rate increased only with small change point probabilities, not more extreme ones (*Fig 3E*). A closer inspection of participant’s behaviour revealed that these unexpected findings may be due to the fact that subjects sometimes lagged slightly behind the optimal model. The task was not self-paced and therefore some delay in participant’s expressed behavioural response is not unexpected, reflecting the time required to process incoming information and translate it into an overt response. Such occasional lags should harm the effect of prior uncertainty less than the effect of change point probability: being fairly smooth from trial to trial (see *Fig 2A*), a strict trial-by-trial alignment between participants and the model is not necessary. In contrast, the change point probability peaks sharply only on a few trials, making trial-by-trial alignment more critical, especially at extreme change point probabilities. To assess whether trial-wise (mis)alignment underestimated the correlation between latent uncertainty factors and learning rate, particularly for change point probability, we implemented an optimization procedure that selectively shifted some subjects’ updates backward by one trial to compensate for a potential delay with respect to the optimal model. For each participant, we searched for the configuration of trial shifts that maximized the correlation between the latent uncertainty factors and the (shifted) apparent learning rate. This procedure improved the correlation with apparent learning rates for each latent uncertainty factor beyond chance level (both *p* < 1 × 10^-13^, see *Methods* and *Supplementary Material, Fig S3*; note that chance level was estimated following the same optimization procedure to avoid circularity in the assessment of the improvement). The improvement was larger for change point probability than prior uncertainty, consistent with our prediction (see *Supplementary Material, Fig S3*). Since lags seem to exist, it could be useful to realign the data before conducting pupillometry analyses. However, below we used the raw data because the majority of trials are not lagged, shifting only some trials would create gaps, and identifying exactly which trials are truly lagged is methodologically impossible (at least with our methods, see *Supplementary Material, Fig S3*).

### Uncertainty factors are encoded within distinct pupil dilatory responses

To examine the association between uncertainty factors and arousal-linked neuromodulation, we ran a multiple linear regression on pupil diameter signals for each peri-stimulus time point in a window defined around an observation (-250 to 1400 ms). The regression model included as predictors prior uncertainty and change point probability, as well as multiple covariates (see *Methods*). The analysis was performed on both phasic and tonic pupil signals, defined as changes relative to a pre-stimulus baseline (-250 to 0 ms) and the absolute (i.e. nonbaseline-corrected) signal, respectively [34] (see *Methods*). We demonstrate separable uncertainty effects between the distinct pupil-dilatory signals: phasic changes in pupil diameter displayed only an effect of change point probability, while the tonic pupil dilatory signal was influenced only by prior uncertainty. The effect of change point probability on the phasic signal emerged 630 ms after the observation presentation until the end of the epoch (*Fig 4B*), which corresponded to a secondary dilatory phase in the baseline-corrected signal conventionally attributed to cognitive processing post orientation [41] (*Fig 4A*). The effect on the tonic signal was evident across the entire duration of the epoch, which importantly included the period prior to observation presentation signifying the presence of accumulative processing arising from past experiences (*Fig 4C*). In other words, the tonic pupil signal reflected prior uncertainty even before observation onset.

**Fig 4.**
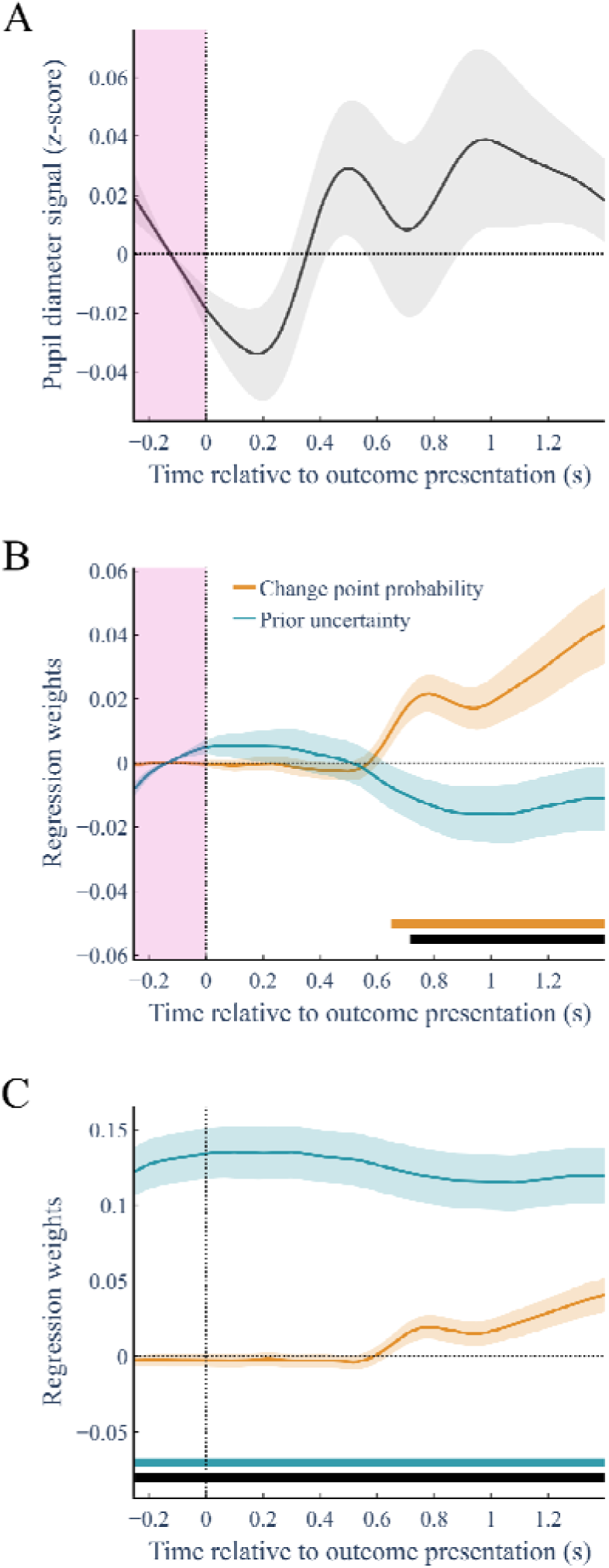
Phasic- and tonic-associated pupil diameter dynamics reflect distinct uncertainty factors. **(A)** Two aspects of pupil responses were evaluated: a phasic response, which was defined by the baseline-corrected (-250 to 0 ms) pupil diameter signal (highlighted on plots), and a tonic response which was defined by the non-corrected, absolute signal. Participants’ displayed transient pupil dilations to outcomes (*i.e.* stimuli within the sequences), as revealed by the average, z-scored baseline-corrected pupil signal. **(B and C)** The effect of uncertainty factors on the change in pupil signals were evaluated using multiple linear regression, along with covariates (see *Methods*). This regression model was estimated across trials separately for each peristimulus time point (-250 to 1400 ms) for both the phasic **(B)** and tonic **(C)** pupil diameter responses. We found an exclusive coupling between pupil signals and uncertainty factors effects: phasic-associated responses displayed an effect on change point probability from 630 ms after observation presentation until the end of the epoch, while tonic-associated responses displayed an effect of prior uncertainty across the entirety of the epoch. Horizontal lines display significant time points for the corresponding (color-coded) uncertainty parameter based on cluster-based permutation analysis (*p* < 0.05, two-tailed, FWE cluster-corrected for multiple comparisons across time). Black horizontal line represents significant time points for a difference between uncertainty parameters based on cluster-based permutation analysis (*p* < 0.05, two-tailed, FWE cluster-corrected for multiple comparisons across time). All plots display the mean ± s.e.m.

To evaluate whether these effects could be attributed to some artefact associated with increased arousal instigated through moving the cursor in response to a new observation, and establish more broadly the generalisability of the observed effects, we ran the same analysis on pupil dilatory responses obtained during a previous study conducted using auditory stimuli (n = 15). The structure of this task was extremely similar to the main study, with one important difference being that participants did not provide an explicit estimate after each observation (see *S1. Appendix* for further task details), but only occasionally on trials excluded from the analysis. The same normative model and regression analysis described in the main study were applied to the observed sequences. The results of the analysis demonstrated the same coupling between pupil signals and uncertainty factors within the auditory probability learning task: phasic changes in pupil diameter displayed an exclusive effect of change point probability and tonic changes were modulated only by prior uncertainty (see *Supplementary Material, Fig S4*). Therefore, the observation that distinct uncertainty factors exert influence on distinct aspects of the pupil response are amodal (not specific to either vision or audition) and linked to the learning process itself, rather than behavioral reports.

### Encoding of uncertainty factors in pupil size modulates participants’ apparent learning rate

We next examined whether identified pupil signatures of latent uncertainty factors were predictive of participants’ apparent learning rate. To achieve this, we extracted the average pupil signal corresponding to each of the observed significant uncertainty factor effects (*Fig 5A*). For the phasic pupil response (*i.e.* baseline corrected) the average was computed from 630 to 1400 ms relative to the observation, where the effect of change point probability was identified. For the tonic pupil effect (*i.e.* non-baseline corrected) the average was taken from the period -250 ms to 0 ms before an observation, where the effect of prior uncertainty was significant. The two average pupil signals were negatively correlated (Pearson’s correlation = -0.45 ± 0.08; ; *t*_(27)_ = -29.84, *p* < 0.0001). When these two quantities were regressed (in the same regression model) on to participants’ apparent learning rate, both the average phasic pupil effect and tonic pupil effect displayed an increasing positive influence (Fig 5B). The phasic and tonic effect sizes were close in magnitude and not significantly different (phasic - beta: 0.12 ± 0.02, tonic - beta: 0.14 ± 0.02; *t*_(27)_ = -1.19, *p* = 0.246; *Fig 5B insert*). However, this observation is still compatible with prior uncertainty exerting a larger effect on the regulation of the apparent learning rate. This is because the similar effect sizes for the pupil signals on learning rates were estimated with normalized (z-scored) changes in tonic and phasic signals, but it should be remembered that changes in tonic pupil size accounted for by prior uncertainty are three times larger than the changes in phasic pupil size accounted for by change point probability (compare effect sizes in *Fig 4B* and *4C)*.

**Fig 5.**
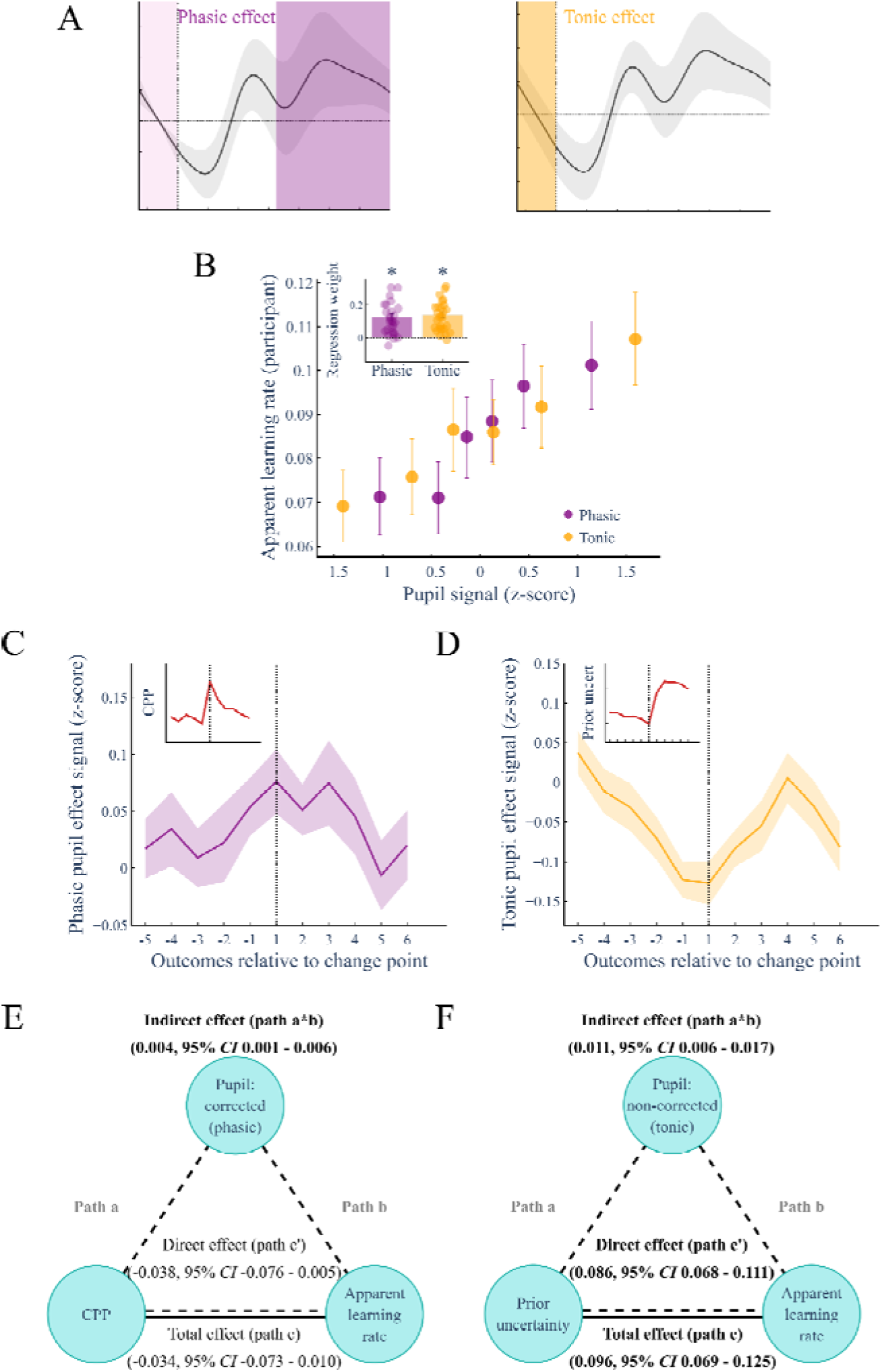
Tonic pupil diameter changes mediate the effect of prior uncertainty on participants’ apparent learning rate. **(A)** The peristimulus regression analysis identified separable effects of change point probability and prior uncertainty within the baseline-corrected and nonbaseline-corrected pupil signals, respectively (see *Fig 4B and 4C*). The traces show the time windows of the observed effects for both signals within which we average the respective pupil signal to obtain one value per trial. **(B)** Tonic and phasic pupil signal effects exert positive influences on participants’ apparent learning rate. To illustrate these effects, the data were binned into 6 equal quantiles based on each pupil signal and averaged within-participants. Analysis was performed using multiple linear regression, the group-average and individual regression weights are displayed in the inset (Phasic: *p* ≈ 1 × 10^-5^ ; Tonic: *p* ≈ 1 × 10^-7^). There was no difference between the effect of the two latent uncertainty-associated variables (*p* = 0.246). **(C)** The tonic pupil signal decreases prior to a change point, before increasing after it; this pattern parallels the prior uncertainty around change points (see inset). **(D)** The phasic pupil signal did not visually display as strong a similarity to change point probability around change points (see inset). **(E)** Given the effect of prior uncertainty on participants’ apparent learning rate, a mediation analysis was performed to examine whether the associated tonic pupil diameter response could, at least in part, explain this effect. The indirect path was significant indicating the tonic-associated pupil responses, at least in part, mediated the effect of prior uncertainty on participants’ apparent learning rate. **(F)** We also demonstrated that an effect of change point probability on participants’ apparent learning rate could be identified when considering the mediating effect of the phasic pupil signal, as indicated by the significant indirect effect. Mediation analyses were performed for each participant and then averaged across-participants, with regression weights and confidence intervals (95% *CI*) based on bootstrapping with 5000 samples. Significance was determined based on the 95% *CI* not spanning across zero, with all instances displayed visually as path texts highlighted in bold. All other traces display the mean ± s.e.m.

We note the change in the tonic pupil signal around change points displayed a qualitative resemblance to the corresponding change in prior uncertainty (*Fig 5D*). Given the dominant effect of prior uncertainty on participants’ apparent learning rate, coupled with the observation that the associated arousal representation of this uncertainty factor in the tonic pupil response was also predictive of apparent learning rates, we then ran a formal mediation analysis of the effect of prior uncertainty (predictor) on learning rate (dependent variable) with the average tonic pupil signal as a mediator (see *Methods* for more information). We demonstrate that the effect of prior uncertainty on participants’ apparent learning rate can, at least in part, be attributed to an effect mediated by tonic pupil responses (*Fig 5F)*. The change in the phasic pupil signal appeared to track change point probability around change points, though visually this was not as apparent as the association between prior uncertainty and the tonic pupil signal (*Fig 5C*). Though we observed a non-significant effect of change point probability on participants’ learning rates, we ran a similar mediation analysis, to examine whether the effects of change point probability (predictor) on learning rate (dependent variable) were carried indirectly through shared associations with phasic pupil responses (mediator). The analysis indicated that change-point probability exerted an influence on participants’ apparent learning rates via this pupil signal (*Fig 5E*).

### The impact of latent uncertainty factors on learning is dependent upon tonic arousal levels

The mediation analyses above indicate that pupil-linked arousal signals carry part of the effect of uncertainty on learning. This raises a complementary question: does the strength of the relationship between latent uncertainty factors and learning rate itself depend on the arousal state in which learning occurs? To address this, we ran the same regression analyses conducted previously to examine how latent uncertainty factors predict participants’ apparent learning rate (see *Fig 2*) after splitting the data into quartiles based on the average tonic pupil signal. Comparisons across tonic arousal quartiles were performed using repeated measures ANOVA (Greenhouse-Geisser adjustments applied where the assumption of sphericity was violated).

As a manipulation check, we first confirmed the uncertainty quantities themselves were appropriately distributed across quartiles. Change-point probability did not differ significantly across tonic pupil quartiles (F_(3, 81)_ = 0.19, p = 0.902), ruling out the possibility that any quartile differences in its effect on learning rate simply reflect differences in the underlying quantity. Prior uncertainty increased significantly across quartiles, as expected given its established relationship with tonic pupil signalling (F_(3, 81)_ = 103.16, p < 0.001); see *Supplementary Material, Table S2)*. Of note, the average phasic pupil signal decreased significantly across tonic quartiles (*F*_(3, 81)_ = 212.53, *p* < 0.001), consistent with the negative tonic-phasic relationship observed across the dataset. We then compared per-participant regression weights relating each uncertainty factor to apparent learning rate across quartiles. The effect of prior uncertainty increased numerically, though we did not see a statistically significant linear effect across quartiles (*F*_(3, 81)_ = 2.44, *p* = 0.070). Importantly, the effect of change-point probability differed significantly across quartiles (*F*_(3, 81)_ = 5.39, *p* = 0.001). Pairwise comparisons (FDR-corrected) confirmed the highest tonic quartile displayed an unexpected negative effect of change-point probability on learning rate, which differed significantly from each of the other three quartiles (Fig 6A).

**Fig 6.**
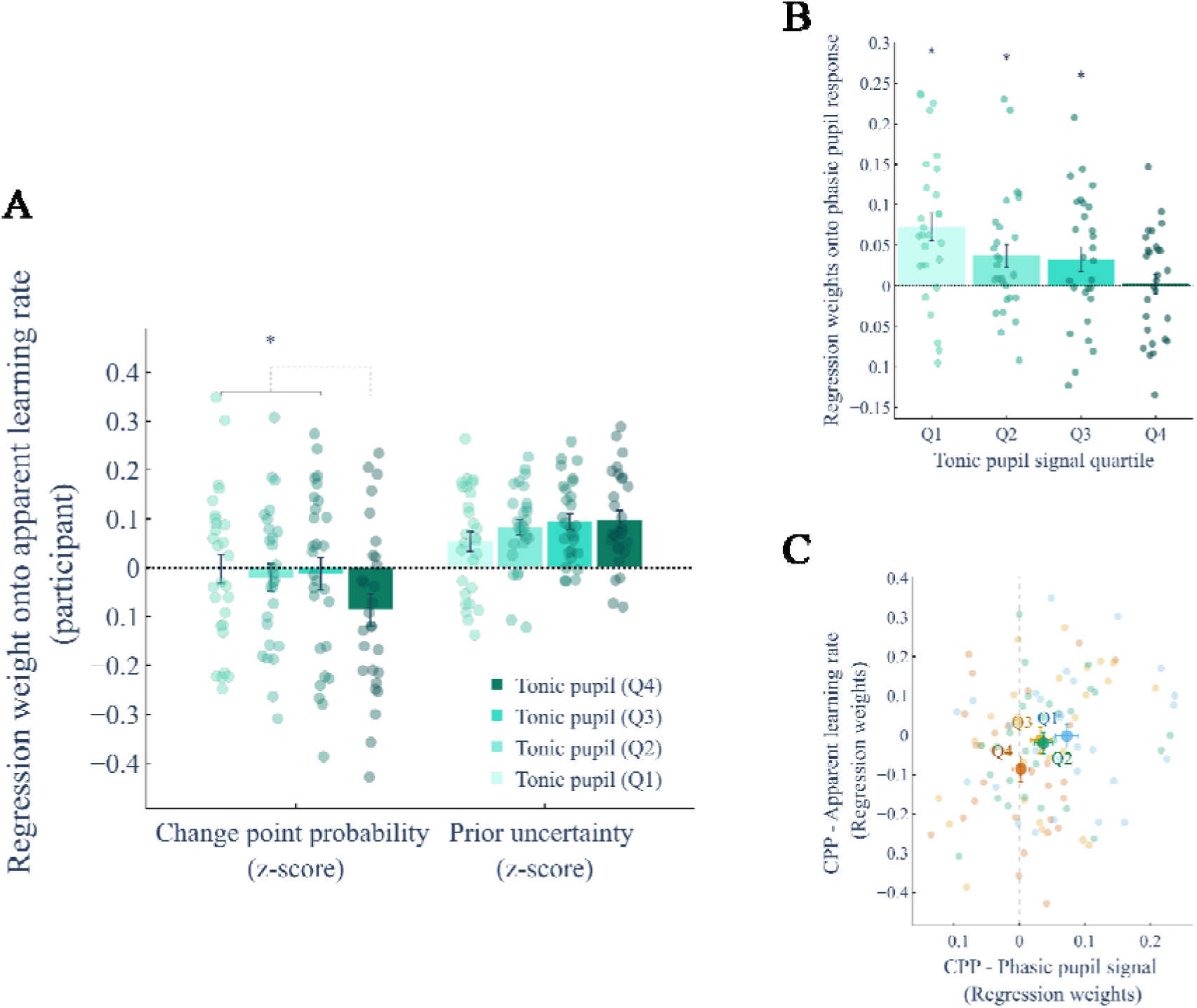
Effect of tonic arousal level on latent uncertainty drivers of participants’ apparent learning rate. **(A)** Change point probability and prior uncertainty (plus defined covariates) were regressed onto participants’ apparent learning rates with the data split into quartiles based on the average tonic pupil signal. Numerically, the effect of prior uncertainty increased across arousal levels, though linear differences were not significant between quartiles. Conversely, the effect of change point probability (CPP) differed between quartiles, with a negative effect observed at the highest tonic pupil level (*: post-hoc FDR-corrected pairwise comparisons, *p* < 0.05). **(B)** To understand the surprising negative effect, the dynamics of the effect of CPP on phasic pupil responses were examined across tonic pupil levels. The regression weights of CPP on phasic pupil responses differed across tonic pupil levels. Compared to zero, significant effects were evident for quartiles 1-3, but was not significant at quartile 4 (*: post-hoc FDR-corrected pairwise comparisons, *p* < 0.05). **(C)** A graphical representation of the change in the association between CPP and both apparent learning rates and phasic pupil response across different tonic pupil levels. The results indicate that tonic arousal levels modulate how uncertainty factors influence apparent learning rates, and show a particular impairment to the encoding of change-point information in phasic arousal signals at higher tonic arousal states.

In light of the mediation results, to better understand this unexpected negative effect of change point probability on apparent learning rate when tonic arousal was high, we tested whether the coupling between change point probability and phasic pupil response itself varied with tonic arousal. We demonstrate that the beta coefficients differed across the different levels of tonic arousal (*F*_(3, 81)_ = 5.94, *p* = 0.001). Within-participant effects of change-point probability on phasic pupil response were significantly positive compared to zero across the first three quartiles (FDR-corrected, all p < 0.05; Average betas = 0.03 - 0.07) but were no longer significant at the highest quartile (beta *= 0.002 ± 0.066,* p = 0.424; *Fig 6B*).

These findings suggest that tonic arousal differentially influences uncertainty-driven learning. Whereas prior uncertainty exerts a qualitatively stronger influence on learning rate under higher tonic arousal, the effect of change-point probability becomes negative at the highest tonic levels. This shift coincides with reduced coupling between change-point probability and phasic pupil responses, consistent with impaired encoding of change-point information in phasic arousal signals (*Fig 6C*). The results indicate that the activity profile of ascending arousal systems play an important role in encoding distinct latent uncertainty factors that shape the integration of new information during probability learning.

## DISCUSSION

This research utilised an inferential task to understand how arousal systems, indexed through pupil dilatory responses, represent latent uncertainty factors and shape human probability learning. Similar to normative theory, we found that uncertainty (or its reciprocal, confidence) in prior beliefs is leveraged to update beliefs following new observations in an environment that makes changes difficult to identify [14]. We also add to a body of accumulating evidence that identify a separation in how latent uncertainty factors underlying statistical learning map onto arousal profiles, with tonic and phasic pupil signals covarying with prior uncertainty and change point probability, respectively [18,34,35,37]. Furthermore, we demonstrate that both of these pupil signals, at least in part, play a role in mediating how latent uncertainty factors are transformed into belief updates during human probability learning. Our results suggest that, within the context of probability learning, belief updates are driven by the representation of separate normative uncertainty factors (*i.e.* prior uncertainty and change point probability) across distinct activation modes of the ascending arousal systems, which interact to appropriately drive learning based on the prevailing uncertainty context.

We relied on a normative modelling framework to characterise how latent uncertainty factors shape learning in a dynamic environment. Consistent with theoretical predictions, probability learning was dominated by prior uncertainty, replicating a previous study [14]. Within this Bayesian formulation, prior uncertainty is epistemic in nature: it reflects an estimate of the lack of confidence in current beliefs about the environment [42]. Our results therefore support the view that adaptive probability learning is principally governed by a confidence-weighting mechanism, whereby belief updates are scaled according to uncertainty about prior expectations [5,13,18,34]. Several lines of evidence support the notion that the brain represents a computationally rational form of confidence. Here, “rational” means it is qualitatively similar to the Bayes optimal solution. This confidence that regulates learning seems accessible to introspection, as model-based confidence correlates with subjective confidence reports [43,44]. Converging neuroimaging evidence has also identified similar neural correlates of confidence signals across multiple tasks [17,44–50]. More broadly, our behavioural results may also suggest that the influence of confidence on learning is context-sensitive, becoming more prominent when new observations are weakly informative, as in probability learning, compared to contexts with higher informational gain, such as when learning about magnitudes [14].

At the mechanistic level, confidence may modulate intrinsic properties of brain states to amplify or dampen belief updates to surprising observations [34]. One mechanism by which this may occur may be arousal state, which subserves the emergence of large-scale patterns in many neural subsystems [28]. Our observed pupil dilatory responses indicate that this confidence-weighting mechanism, as well as the perception of change points, is dependent upon arousal state and neuromodulation. In line with previous research, we demonstrate that the representation of uncertainty factors during probability learning adheres to a separation of prior uncertainty and change point probability between tonic and phasic arousal responses, respectively [18,35,37]. We confirm this separation within the context of probability learning [34], and show that both pupil responses carry information that determines learning. This separation was not an artefact induced through participants’ reports as confirmed by similar effects observed in our supplementary analysis of an audio probability inference task requiring no overt response. Previous work has shown that both change-point probability and relative uncertainty (*i.e.* prior uncertainty) influence apparent learning rates through shared variance expressed in the P300 EEG component [15,16]. The P300 component has been proposed as the cortical index of a broader orienting response to novel and uncertain events, consistent with its covariation with phasic autonomic measures such as pupil dilation and skin conductance [51]. More precisely in computational terms, the P300 may index the updating process itself [52,53], and therefore recapitulate the multiple factors that influence learning. In contrast, our findings show that tonic and phasic arousal states do not reflect the update itself, but the dissociable uncertainty factors that regulate belief updating, by selectively tracking prior uncertainty and change point probability, respectively.

Our results align with the findings of Nassar and colleagues (2012), who showed in a magnitude inference task that outcome-evoked pupil dilation over a period of 0-2 s after feedback encodes change-point probability, whereas mean pupil size over the same period reflects relative uncertainty. In their study, outcome-evoked pupil responses predicted participants’ learning rate, whereas average pupil size was a non-significant predictor. Here, we extend these findings, identifying pupil-linked arousal as an important regulator of apparent learning rates within the context of probability learning. We demonstrate that in probability learning, tonic pupil responses are predictive of adopted learning rates. The stronger contribution of this tonic component — relative to the non-significant effect of average pupil size reported by Nassar *et al.*, (2012) — may reflect the greater importance of prior uncertainty in probability learning compared with magnitude learning [14]. We also provide evidence suggesting an intricate interplay between tonic and phasic arousal states in determining how latent uncertainty factors drive learning. The findings together suggest effective regulation of learning, be that probabilities or magnitudes, would appear to be dependent upon fine tuning the balance between tonic versus phasic activation of arousal systems in a way that is appropriate for a given context.

Non-luminance-dependent pupil dilations, reflecting global arousal, are widely used as an indirect index of locus coeruleus (LC) activity and noradrenaline (NA) release [25,54]. The strongest evidence for this link comes from studies showing simultaneous activation of LC firing and pupil responses and demonstrations that LC microstimulation evokes pupil dilation [24,26,55,56]. In humans, pupil diameter covaries with LC BOLD activity both at rest and during cognitive tasks [57]. Although non-luminance-dependent pupil dilations are not exclusively controlled by the LC-NA system [24,26,58,59] and are state dependent [60], our results can speculatively be interpreted in relation with LC activity, which we now discuss.

Several theoretical accounts propose that the LC–NA system plays a central role in behavioural adaptation, particularly to environmental change [9,61–64]. Supporting a broad interpretation of these perspectives, LC activity tracks changes in environmental contingencies and often precedes behavioural adjustments [65–67], while causal manipulations of the NA system — including LC optogenetic activation, prefrontal NA deafferentation, and NA-targeted pharmacological interventions — modulate flexible learning and choice variability [66,68–74]. Our finding that participants’ apparent learning rates were predicted by both tonic and phasic signals is compatible with the hypothesis that the brain may exploit distinct LC activity modes to adapt learning to environmental context. In the probability learning task used here, the dominance of prior uncertainty in shaping learning rates and the association between high tonic arousal with altered phasic representation of change point probability aligns with adaptive gain theory [75]. Within this framework, tonic LC activity modulates the selectivity of sensory processing: increased tonic activity is described as a “distractible” mode, defining a broad attentional state based on a global increase in neural gain where new information can more effectively challenge current beliefs. Consistent with this account, larger baseline pupil responses are predictive of individuals engaging in favouring exploration over exploitation in rewarded behavioural tasks [76–78]. Tonic pupil size also correlates with a broad attentional focus and these pupil changes are predictive of the strength of fMRI responses, as would be predicted by a global modulation of neural gain [79]. Our analysis across increasing levels of tonic arousal suggests that the highest tonic states may exceed the range in which phasic signals effectively track environmental change. Under these conditions, the coupling between phasic pupil responses and change-point probability weakened, accompanied by a reduced contribution of change-point probability to learning. This pattern is consistent with a reconfiguration of uncertainty processing under elevated tonic arousal. Accordingly, during probability learning driven by prior uncertainty, a high tonic arousal state may be preferable, especially after change points, as information is sought from new observations limited in their informativeness.

The LC also plays an important role in memory [80–82]. Phasic pupil responses have been shown to demarcate episodic memories [83]. This is compatible with the demonstration that the observed phasic pupil response is sensitive to the probability of changes in the generated sequence. Yet, higher tonic LC activity ameliorates phasic dilation effects at event boundaries, which may hold implications for episodic memory formation [83]. High tonic arousal may change how information is processed by engaging higher-level transmodal regions, including the hippocampus [84], which may emphasise the importance of functions such as memory in the interpretation of the salience of new information. Here, a high tonic arousal state due to increased uncertainty in prior beliefs may require such engagement to track relationships between clusters of observations in order to build confidence in inferences concerning the underlying statistical structure of the environment. Our analysis across tonic arousal states is compatible with this interpretation, showing that at the highest tonic levels the phasic representation of change-point probability becomes attenuated while learning remains strongly influenced by prior uncertainty. This pattern suggests an interaction between the dominant uncertainty factor driving learning and the manner in which uncertainty is represented. Here, elevated tonic arousal may preferentially support the maintenance and refinement of belief confidence over the signalling of environmental change. To this end, the activity profile of the LC may provide the gating mechanism defining precisely how the brain appropriately configures to perform predictive inference in changing environments.

We now discuss the choice of model used in our analysis. We used a hidden Markov implementation, also known as Bayesian filtering, of an ideal Bayesian observer [34] which assumes the brain computes the full posterior over possible outcomes. In a previous study [85], this model was reported to best fit behavior in another dataset using the present probability learning task. The benchmark included a range of alternative models — both approximately Bayesian (*e.g.* hierarchical Gaussian filter, volatile Kalman filter, adaptive mixture of delta rules, reduced Bayesian, and change point models) and clearly non-Bayesian (*e.g.* proportional-integral-derivative controllers, adaptive delta rules). More precisely, this previous study used a parameterized ideal observer model, in which the volatility assumed by the model was subject-specific, rather than matching the generative value. We explored this possibility here but it did not improve the results substantially. This computationally demanding form of learning required by the Bayesian model poses the question of whether its accuracy is worth the cost. Across tasks, recent evidence suggests that higher-level systems may arbitrate over multiple algorithmic implementations of increasing complexity, dictating which controller drives behaviour given the complexity of the task [86]. There appears a small window across stochastic, low volatility environments, where the computational cost of the Bayesian model is worth the added accuracy that past observations offer in terms of future predictions [87]. In the present task, the generative volatility was 0.05 (one change point every 20 observations, on average), which resides within a space that could benefit from using a Bayesian model [87]. But, this in turn raises the longstanding question of the biological feasibility of Bayesian inference [88–90]. Our lab has recently provided evidence that an artificial neural network displays the capacity to solve probability learning tasks quasi-optimally with a very small number of units (about 3 to 10), arguing in favor of the biological plausibility of such computations, but only when the network had a gated, recurrent neural architecture [91]. LC activity may provide a gating mechanism in brain circuits, akin to the one of artificial recurrent gated neural networks, that enables confidence-weighted learning.

We note that the tonic pupil response only partially mediated the effect of prior uncertainty on participants’ apparent learning rates, indicating that additional factors contributed to either pupil dynamics or learning behavior. Our mediation analysis may not capture the nonlinear coupling of arousal to brain states dynamics [28] which may underlie learning. Moreover, areas such as the anterior cingulate cortex (ACC) have been implicated in the updating of internal models and control of learning rate [3,33]. Top-down processing, including from the ACC, influences pupil-linked-arousal [24,92], but may also have other effects on learning, which would result in the tonic pupil size mediating (in the statistical sense) only partly the effect of prior uncertainty onto the apparent learning rate. Regarding pupil measurements, several limitations should be considered. Pupil diameter can be affected by interactions with other autonomic processes. Recent evidence shows that pupil constriction and dilation correspond to inspiratory and expiratory phases of breathing, respectively [93], which may confound interpretations of pupil responses to latent uncertainty. Additionally, our protocol does not allow the identification of any potential strategies participants may have employed to perform the task [39], which may have required more or less mental effort that may also be captured within pupil dilatory responses [94]. A further limitation may concern our definition of the tonic pupil response. In the present study, tonic and phasic pupil signals were operationalized as non-baseline and baseline corrected dilatory responses, respectively. With short interstimulus intervals (here, 1.5 s), this approach may inflate tonic estimates due residual phasic responses from previous trials. Emerging algorithms may offer improved decomposition of tonic and phasic components and circumvent this limitation [95]. We employed this new technique to check our data and confirm that our results hold when adopting the deconvolution method (*Supplementary Material, Fig S5*).

In summary, our results support human probability learning to be based, at the computational level, on a qualitatively optimal integration of latent uncertainty factors, with confidence in prior beliefs serving as the primary driver of belief updating. Importantly, we show that uncertainty-guided regulation of probability learning is implemented via different arousal-related responses. Interactions between arousal system activations suggest neuromodulation dynamically regulates which uncertainty signals are prioritised and used to guide belief updating in continuously changing environments.

## METHODS

### Code and data availability

The collected data, code for the experiment, and the analysis code are all available on the following GitHub repository: https://github.com/TheComputationalBrain/Adalearn/tree/main/Project/adalearn_pupil

### Participants

Thirty-two participants were recruited via *the Paris Brain Institute PRISME Core Facility (<u>RRID:SCR_026394</u>), Paris, France.* They provided written informed consent prior to participation. The study was approved by the Ethics Committee of the Paris-Saclay University (Comité d’Ethique de la Recherche; reference: CER-Paris-Saclay-2023-010) and took place in the PRISME behavioural facility. To evaluate participants’ proficiency with the task, we used the correlation coefficient between participants and ideal observer model estimates as a performance criterion. A negative relationship was taken as evidence that the participant failed to fully comprehend the task instructions. One participant displayed a negative correlation (Pearson’s correlation = -0.16) and was therefore removed from all further analysis. For the remaining 31 participants (17 female), the average age was (mean ± standard deviation) 38 ± 11 years, ranging between 19 and 63 years. Participants spent 4 ± 3 years in higher education, and 8 reported additional mathematical training as part of their education or occupation. All participants reported normal, non-corrected vision. To ensure the quality of the analysed pupil signal, checks on the degree of missing data led to the exclusion of 2 participants from analyses involving pupil diameter (further explained in *Pupil diameter recording and preprocessing*). A further participant was removed from all pupil analysis due to a failure in the recording of the pupil data. Accordingly, 28 participants were included in the pupil-based analyses, while all other analyses were performed on 31 participants. Participants received 25€ for participation, in addition to a performance-based bonus up to 5€.

### Task

The probability learning task has previously been described [14]. One hundred and fifty sequences were generated for the task [14], which were randomly sampled without replacement from these sets. The generation of the sequences was based on a Bernoulli process, whose parameter (*i.e.* the hidden generative probability) changed with a probability of zero for the first six observations following a change point and 1/20 for all trials thereafter. As previously described, the generative probability was sampled uniformly between 0.1 and 0.9 both initially and after a change, with the latter subject to the constraint that the resulting change in the odds-ratio be no less than fourfold. The average number of change points per sequence was (mean ± *SD*) 3.07 ± 0.26, with 18.68 ± 11.31 trials separating each change point.

Task sessions required participants to continuously estimate the hidden generative probability (*i.e.* proportion of blue on the fictitious roulette wheel) from a sequence of 75 observations, by moving the slider along the track in response to each observation. The interval between the onset of each new observation was 1.5 s, with stimulus offset occurring at 1.3 s. Within this period, participants could move the slider using a trackpad to indicate their new estimate. The slider was highlighted to indicate when participants’ estimates were recorded, reflecting its position on the slide tracker at 1.5 s post-stimulus. At the end of each sequence participants received feedback, used to maximise engagement with the task. The feedback displayed participants’ performance on that sequence (quantified as a percentage score), performance-based monetary gain (which was proportional to their score), and the number of change points within the sequence. The number of change points were represented by horizontal lines for each new generative probability. On the horizontal lines, the real generative probabilities and participants’ own estimates were displayed. Participants’ score for a sequence was calculated based on their mean absolute error, evaluated in relation to two reference points: 1) the ideal observer model estimates (high score), and 2) a ‘stay still’ strategy (low score). These raw scores were then transformed with a softplus function to place them on a smooth, positive scale, making it easier to compare participants’ performance relative to these benchmarks.

The task comprised two steps: 1) instructions and training, 2) task sessions. In step one, participants read through the provided instructions before completing a single sequence of the task. Verbal assessment of understanding identified if a further sequence was required for full comprehension. In step two, participants completed 22 task sessions of the probability learning task (*Fig 1*), split into two equal blocks of 11 sequences. The task sessions took between 45-55 mins to complete.

Several modifications were made to the original design to improve its compatibility with pupil diameter recordings. The visual angle of the slider was set to 3° from the centre of the monitor based on a typical sitting distance (approximately 60 cm). This enabled the whole of the slider to be seen by the participant while they maintained a fixed gaze on the presented stimuli, limiting large saccades during the task and any potential distortion of the pupil diameter recording [96]. A semi-circular slider track was preferred to the original horizontal track to facilitate the smaller visual angle and maintain the same distance to all points along the edge of the track from the middle of the monitor in the eventuality of saccades (*Fig 1*). The horizontal position of the slider was mapped onto the semi-circular track, moving from ‘0’ (100% yellow) to ‘1’ (100% blue) from left to right, respectively. The segments of the slider track between “0–10%” and “90–100%” were filled in order to indicate that the hidden probability never lies within these intervals. Each observation was presented in the middle of the slider track with a fixation target overlaid. Blue and yellow colours were maintained in the task, but hues were adjusted in order to match luminosity, calibrated using a photometer and correcting for the individual systems’ gamma curve (colours presented in *Fig 1* do not accurately reflect the colours presented in the task and are displayed as such simply to emphasise the two different outcomes). The task was presented on a homogenous grey background.

### Behavioural analysis

The apparent learning rate in response to each new observation (□*_t_*) was quantified using a delta rule [5,13] based on the degree estimate beliefs (*p*) were updated (*p_t_ - p_t-1_*) in proportion to level of prediction error (*x_t_ - p_t-1_*):

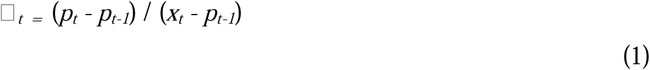

In keeping with previous studies [14,18,20], outliers arising occasionally due to prediction error being very close to 0 were removed from the analysis. Cut-off thresholds of 1.3 and -0.6 were adopted following a previous study [14].

### Normative model

The ideal Bayesian observer model applied to each sequence is described previously [14,34] and is openly available at the following repository: https://github.com/TheComputationalBrain/TransitionProbModel.

Briefly, the model infers the posterior distribution of the hidden generative probability (*h_t_*) from past observations (*x_1:t_*) using a Markovian update rule governed by volatility (ν). It assumes knowledge of the true volatility level and that, when a change occurs, the new generative probability is sampled uniformly from a proposal distribution, *p_0_*(θ), within designated bounds [0.1, 0.9]:

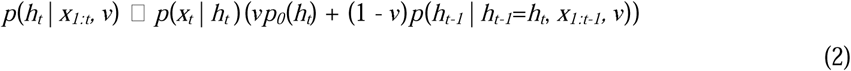

To compute this posterior numerically, the model discretises the probability space over a grid with a resolution of 20.

This model provides a normative account of the optimal solution to the inference problem, and enables the extraction of observation-by-observation changes in the model-derived apparent learning rate and latent uncertainty factors. Two factors were specifically examined: 1) prior uncertainty, the uncertainty of participants about the latent probability before seeing the new observation (also referred to as epistemic uncertainty, or inferential uncertainty Meyniel, Schluneger & Dehaene 2015), which we quantified as the standard deviation of prior probability distribution:

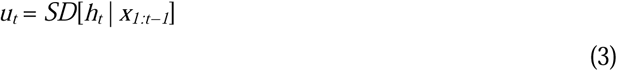

2) change point probability, the probability a change in the hidden state has occurred since the last observation, which is associated with the degree of divergence between the likelihood of new evidence and previous beliefs:

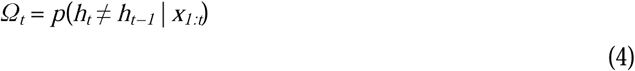

### Lag optimisation analysis

To assess whether belief updating lagged behind the influence of latent uncertainty factors, we implemented an optimisation procedure that evaluated whether shifting trial-wise behavioural estimates improved their correlation. Within each sequence, estimates at each trial were permitted to take the value of the immediately following trial (excluding the final trial of the sequence), thereby modelling a one-trial delay in the behavioural response. To prevent double-counting of information, when a trial’s estimate was reassigned to a preceding trial, the original estimate was removed unless it was itself replaced by an estimate from a later trial. The optimisation algorithm proceeded through ten iterations, during which the shift status of each movable trial was tested in random order. A trial’s update was temporarily replaced by that of the next trial, and the change was retained only if it increased the overall correlation between the participant learning rate and a given uncertainty factor; otherwise, the shift was reverted. The procedure converged upon a stable trial-shift pattern that maximised this correlation. To evaluate whether the observed improvement could arise by chance, a null distribution of optimised correlations was generated for each participant. For this, the participant’s update values were held constant, while the latent uncertainty factor and error terms were drawn from other participants (n = 30). Each null pairing underwent the same ten-iteration optimisation procedure, providing an empirical distribution of chance-level correlations. The difference between the participant’s optimised correlation and the mean of the null distribution quantified the strength of the lag-adjusted association beyond chance. This approach ensured that any apparent temporal alignment between the behavioural and latent computational variables could not be attributed to random correlations or to artefacts of the optimisation process itself.

### Pupil diameter recording and preprocessing

Pupil diameter signals were recorded from both eyes using a Tobii Pro Nano (n = 24) or Tobii Pro Fusion (n = 4) at a sampling rate of 60 Hz. The eyetracker was positioned approximately 10-15 cm under the monitor. The onset of observations were integrated into pupil signal recordings through the use of a websocket connection, facilitating time-locked pupil diameter analysis of each new observation. Our analysis was performed on a nominated eye (*i.e.* the right eye). We checked whether the results held when using the eye with the least missing data for each participant and confirmed that it did, so proceeded with analysis performed on a nominated eye. Preprocessing of the pupil diameter signal was performed using a custom script. Blinks were delineated (adding a margin of 100 ms before and after) with the data linearly interpolated in between. Pupil signals were filtered using a low-pass, bi-directional, 4th-order Butterworth filter (5 Hz), before being epoched within -250 to 1400 ms around each new observation onset. Two pupil diameter signals were evaluated: a baseline-corrected signal, where the mean signal between -250 ms to 0 ms before observation onset was subtracted from each point within an epoch, and a nonbaseline-corrected signal. The baseline-corrected and nonbaseline-corrected signal reflect the phasic and tonic properties of pupil diameter dynamics, respectively. Pupil signals within epochs were z-scored per participant.

Epochs with more than 20% interpolated data were excluded from the analysis. For further quality control, participants with >50% of their total epochs removed due to excessive interpolation were also excluded from the subsequent pupil analysis [97]. This threshold initially flagged four participants. Upon review, we identified that two of the four participants only marginally exceeded this imposed threshold (50.4% and 52.4%, respectively). Given the large number of epochs still available for these participants and in order to prioritise sample retention, it was decided to keep these participants in the group-level pupil analysis. The two participants removed from the analysis recorded a much larger percentage of epochs removed (61.2% and 75.6%, respectively). The two participants who marginally exceeded the threshold were not identified as clear outliers (z-scores: 1.41 and 1.52), unlike the other two participants (z-scores: 2.00 and 2.72). Following this procedure, the total proportion of epochs removed from the dataset used in the pupil analysis was 20.8%. It is important to note, all results hold when the exclusion criteria is strictly adhered to.

### Regression Analyses

All analyses were performed using Python, unless stated otherwise. All regression based models included constants and z-scored predictors and were implemented using ordinary least squares models within the **statsmodels** package.

Prediction of participants’ apparent learning rate based on latent uncertainty factors was performed using a model that included change point probability and prior uncertainty as primary predictors and the interaction term between these uncertainty factors as a covariate.

For the epoch-based analysis of the pupil diameter signal, several covariates were included within the regression model (*equation 5*) to account for sensory, task-related, and behavioural influences on pupil diameter. Observation outcome (blue vs. yellow) was included to control for any residual luminance differences between stimuli. Observation position within the current sequence was included to account for learning and adaptation effects across a sequence, with both squared and square-root transformed terms added to capture potential non-linear changes over time. Task session number was included to account for between-session effects such as practice, habituation, or fatigue. Previous prior belief was included to control for the influence of absolute belief estimates independent of uncertainty. Entropy of the prediction was included to account for uncertainty associated with the overall dispersion of the belief distribution, allowing the unique contributions of prior uncertainty and change-point probability to be estimated. Finally, pupil position, computed as the Euclidean distance from the fixation point, was included to account for gaze-position-dependent variation in pupil size estimation.

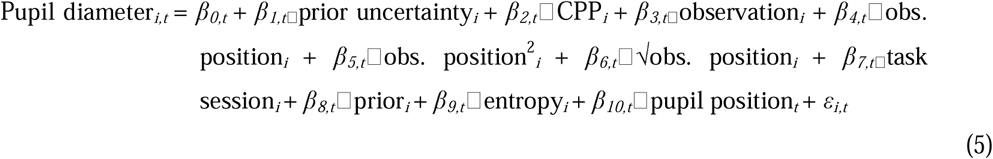

To control for multiple comparisons, the epoch-based analysis evaluated the statistical significance of each latent uncertainty regression weight using a cluster-based permutation analysis (**MNE** statistics package). A one-sample permutation cluster test was performed on the participant × time data matrix with 1,000 permutations, testing against zero under a two-tailed null hypothesis, with alpha set at 0.05. Of note, the same correction procedure was used for the analysis of differences between uncertainty factors within epochs, as well as learning rate dynamics around change point, testing apparent learning rates against baseline means.

For the model assessing participants’ apparent learning rate as a function of identified pupil effects, averaged phasic and tonic pupil responses within time-windows extracted from the epoch analysis were the primary predictors. Covariates included the observation (*i.e.* blue vs. yellow), the position of the observation in the current sequence (as well as the square and square root of this position), the task session number and linear and non-linear learning effects described previously. The moderating effect of the average divergence of the pupil from the fixation point on the pupil diameter response was also entered as a covariate.

The absence of effects between latent uncertainty factors (*e.g.* change point probability) and apparent learning rates did not preclude mediation analysis [98,99], with mediation defined as the presence of an indirect effect. For the mediation analyses, we ran participant-level models to examine whether the effect of latent uncertainty factors (X) on the apparent learning rate (Y) was mediated by associated pupil signals (M). The pupil signals reflected the average signal over a window where the respective uncertainty factor exerted influence (*Fig 5A*). The indirect effect was assessed at the group level using bootstrap procedures. Mediation was implemented using linear regression equations via the **Pingouin** package:

*Path a (X → M):*

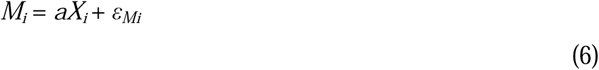

*Path b and direct effect, c′* (*M, X → Y*):

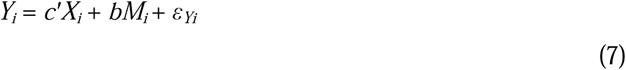

*Total effect (Path c)*:

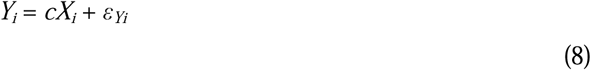

*Indirect effect (Path ab)*:

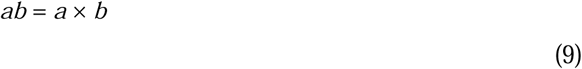

Here, the coefficients represent:

*a*: effect of prior uncertainty on the tonic pupil response;

*b*: effect of the tonic pupil response on learning rate, controlling for prior uncertainty;

*c*′: direct effect of prior uncertainty on learning rate, controlling for the tonic pupil response;

*c*: total effect of prior uncertainty on learning rate;

*ab*: indirect effect of prior uncertainty on learning rate through the tonic pupil response;

At the group level, the significance of the pathways were assessed by constructing bootstrapped 95% confidence intervals across 5000 samples.

## Acknowledgements

The authors would like to extend their thanks to Hamza Sabek and Thaïs Marques for their help in the data collection and piloting the study, respectively.

---

**Fig S1.**
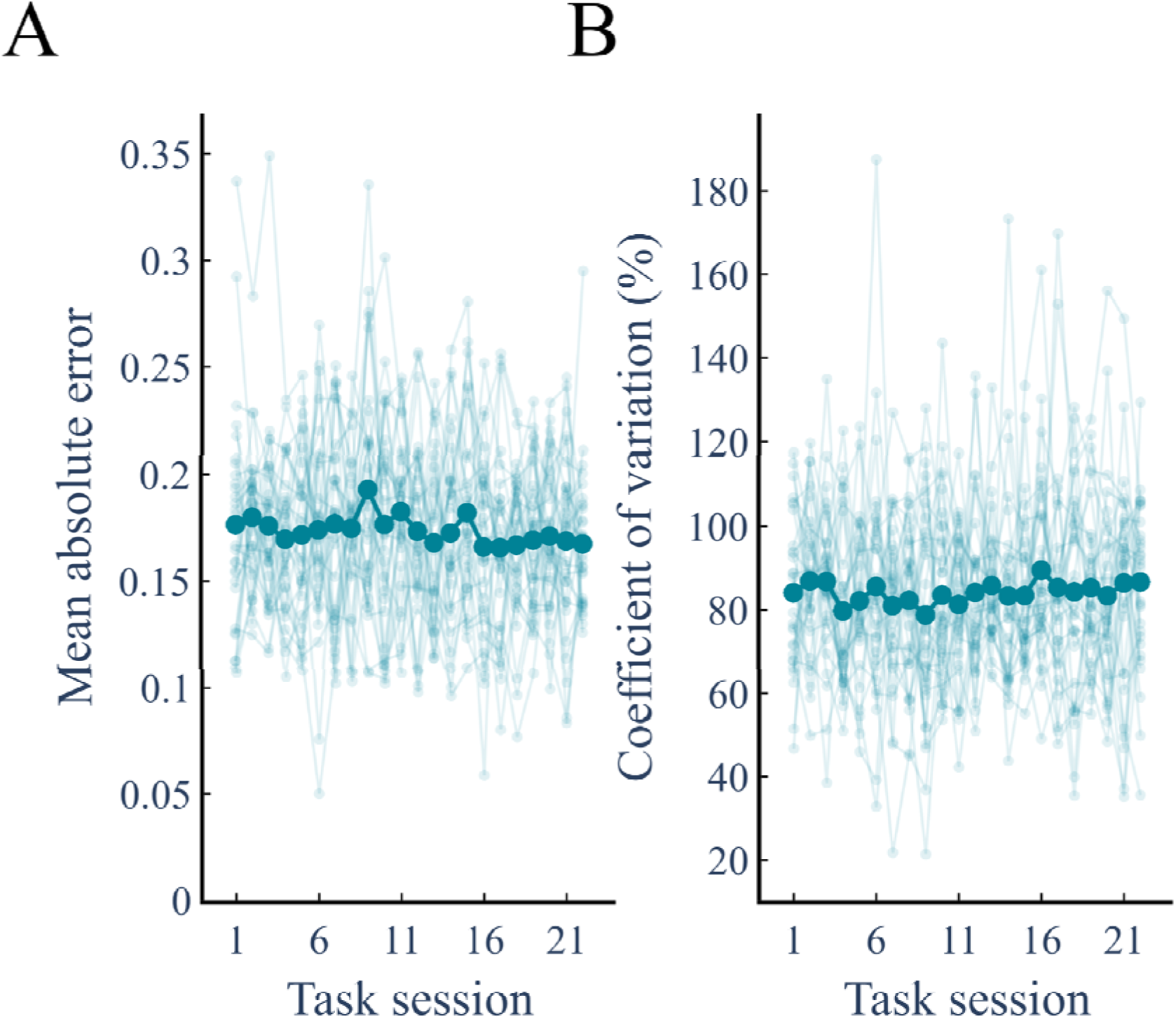
Absolute error and variability in error across task sessions. Participants’ performance was stable over the performed task sessions, as quantified through the **(A)** absolute error (*F* = 0.950, *p* = 0.520) and **(B)** the coefficient of variation (*F* = 0.506, *p* = 0.969). Absolute error was defined by the mean absolute difference between the participant’s estimate and the true value of the hidden generative probability. The variability of this error within each task session was quantified by the coefficient of variation, reflecting the standard deviation in the absolute error for a given task session divided by the mean of that error. Thin dots and lines connecting them each denote one participant, with the larger circles representing the mean across participants.

**Fig S2.**
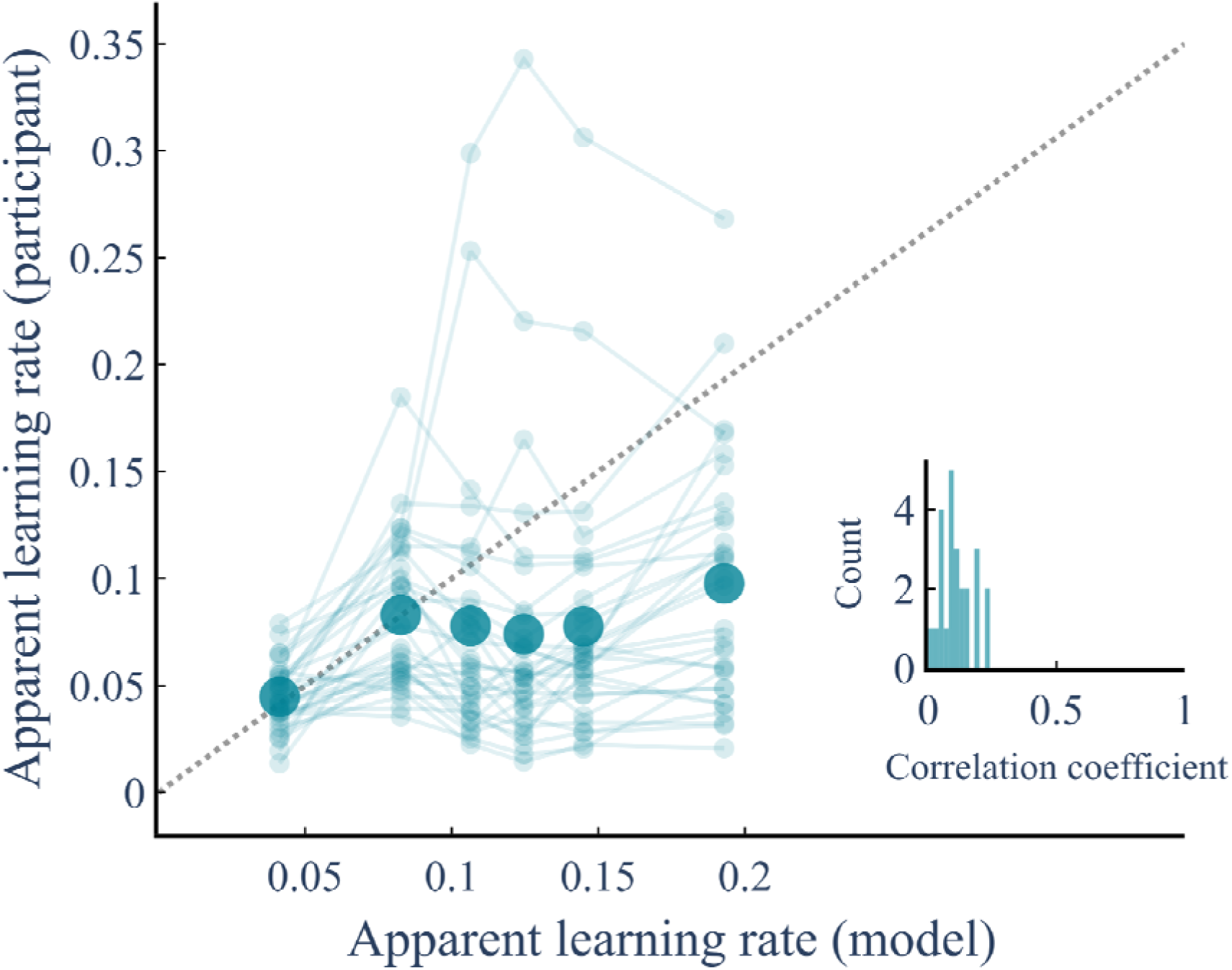
Comparison of participants’ apparent learning rates to model-derived estimates. The data were binned into 6 equal quantiles of model learning rate estimates and averaged within-participants. Small points and lines are the average for each participant, with the larger points reflecting the mean. The dotted line represents the identity line. The insert displays a histogram of the observed correlation coefficients across all participants.

**Fig S3.**
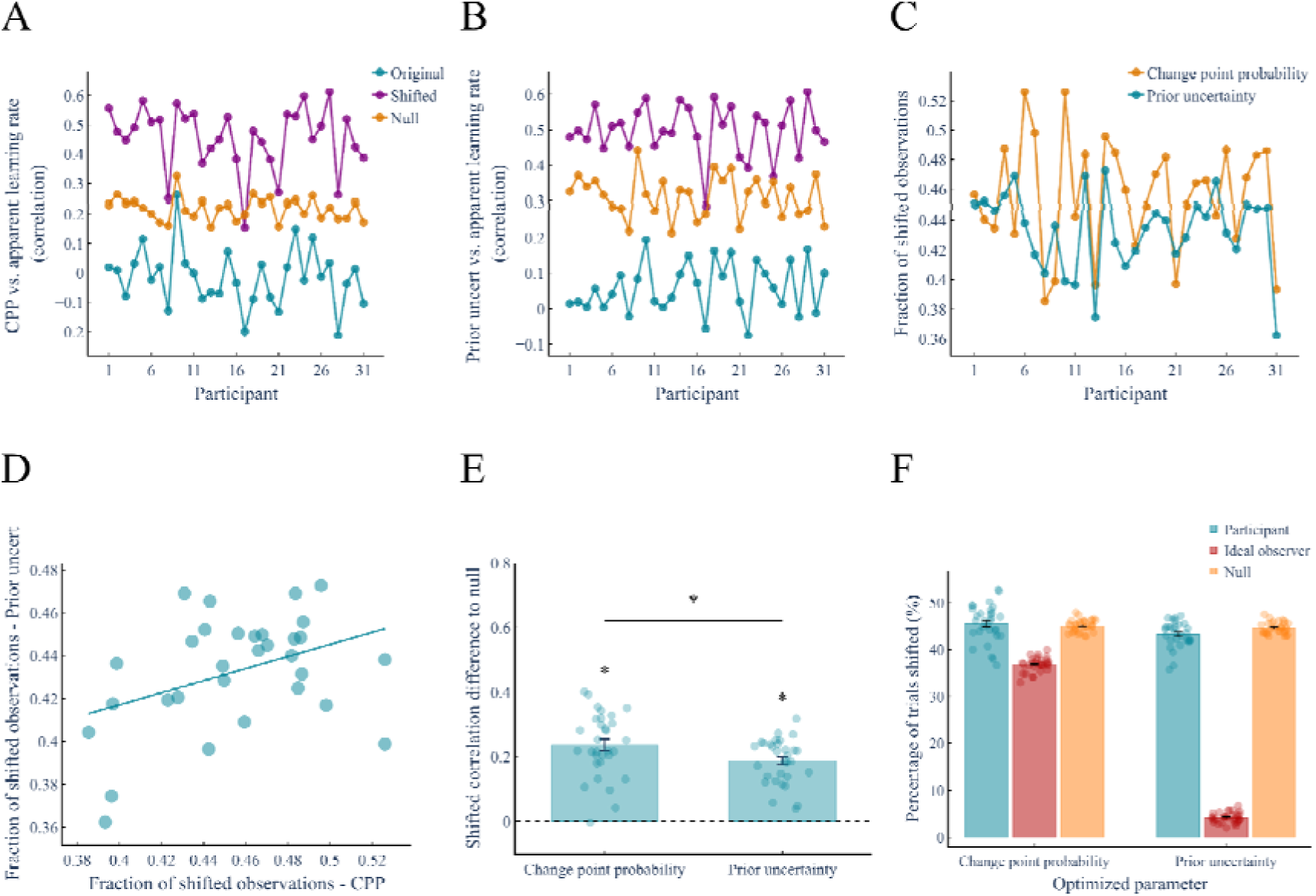
Outcome of “lag” optimization of the latent uncertainty-associated factors relationship with participants’ apparent learning rate. **(A and B)** The optimisation procedure improved the correlation compared to the original correlation and that which may be expected by chance for both change point probability (CPP) and prior uncertainty (mean ± s.e.m; CPP - original correlation: -0.02 ± 0.02, shifted correlation: 0.45 ± 0.02, null correlation: 0.22 ± 0.01, shift vs. null: *t*_(30)_ = 13.33, *p* ≈ 3.90 × 10^-14^; Prior uncertainty - original correlation: 0.06 ± 0.01, shifted correlation: 0.50 ± 0.01, null correlation: 0.31 ± 0.01, shift vs. null: *t*_(30)_ = 14.05, *p* ≈ 1.90 × 10^-15^). **(C)** Between participants, there was a positive correlation between the fraction of trials shifted when optimising the correlation for change point probability and prior uncertainty (Pearson’s correlation = 0.394, *p* = 0.028). **(D)** For each participant, there was a high degree of agreement in the fraction of observations shifted to optimize the correlations. **(E)** The improvement in correlation coefficients is expressed relative to the generated null distribution. The improvement was significant for both quantities **(A and B)**. There was a significant difference between uncertainty factors, with a larger improvement seen for change point probability compared to prior uncertainty (mean ± s.e.m; CPP - 0.24 ± 0.02, Prior uncertainty - 0.19 ± 0.01; *t*_(30)_ = 3.17, *p* = 0.003). **(F)** Proportion of shifted trials returned by the lag optimization procedure, in the participants’ data, the null distribution, and the ideal observer for each optimized parameter. These proportions are similarly high in the participants and the null distribution. For change point probability, it is also high in the ideal observer (where we know the ground truth: 0% of shifted trials). These high proportions are the consequence of the optimization procedure itself, which overestimates the true proportion of shifted trials; they should not be interpreted in absolute terms.

## S1 Appendix.

Auditory probability learning task To evaluate whether movement artefacts contributed to the observed uncertainty-associated pupil dynamics, we checked if the same effects were present through reanalysis of data previously collected in our lab that used an auditory probability learning task very similar to the main task. In brief, the auditory probability learning task consisted of auditory sequences of 420 stimuli, with stimuli composed of 2 clearly distinguishable tones. The separable tones refer to sets of chords (350, 700 and 1400 Hz vs. 500, 1000 and 2000 Hz) played into both ears for 50 ms (including 7 ms rise and fall times). Auditory stimuli were presented with an inter-stimuli interval of ∼1.3 s. On each trial, the probability of a tone was based on a changing Bernoull process. The probability of a change in the underlying generative probability was constant and slightly smaller than the main task (1/75). When a change occurred the generative probability was sampled from the same interval as the main task (0.1- 0.9), with similar constraints employed as the main task (*i.e.* the change in odds ratio should be at least 4-fold). This meant that the task did not have overly long stable sequences and that changes were not so small as to be imperceivable. Importantly, movements during the auditory probability learning task were minimal. Every 22 tones (± 3 tones), participants were asked to rate the generative probability as well as their confidence in their estimate using a mouse-tracker. This was the only movement involved in the task, therefore reducing motor involvement considerably compared to the main task. Participants were presented with at least 4 sequences each, resulting in a total of 1680 observations. Here, we only analysed participants with pupil data which exceeded a quality control threshold related to the amount of missing data within the pupil recording. In keeping with the main task, epoch windows were created around each tone presentation (-250 to 1200 ms). Blinks were accounted for within an epoch through linear interpolation. If the amount of interpolated data exceeded 20% of all data points within that epoch, the epoch was removed from the analysis. If a participant had more than 50% of their total epochs removed they were removed from the analysis (see *Methods*). In total, 15 participants were included in the analysis. The effects of latent uncertainty-factors (*i.e.* prior uncertainty and change point probability) were assessed using a multiple linear regression across each peri-stimulus time-point, in line with the cluster-based permutation analysis performed in the main study. Of note, the two tasks differed in the eyetracker used to record pupil size (in the auditory probability learning task, EyeLink 1000), with the sampling rate of the pupil signal higher in the auditory probability task (500 Hz) compared to the main study (60 Hz). Results of the analysis are presented in *Fig S4*.

**Fig S4.**
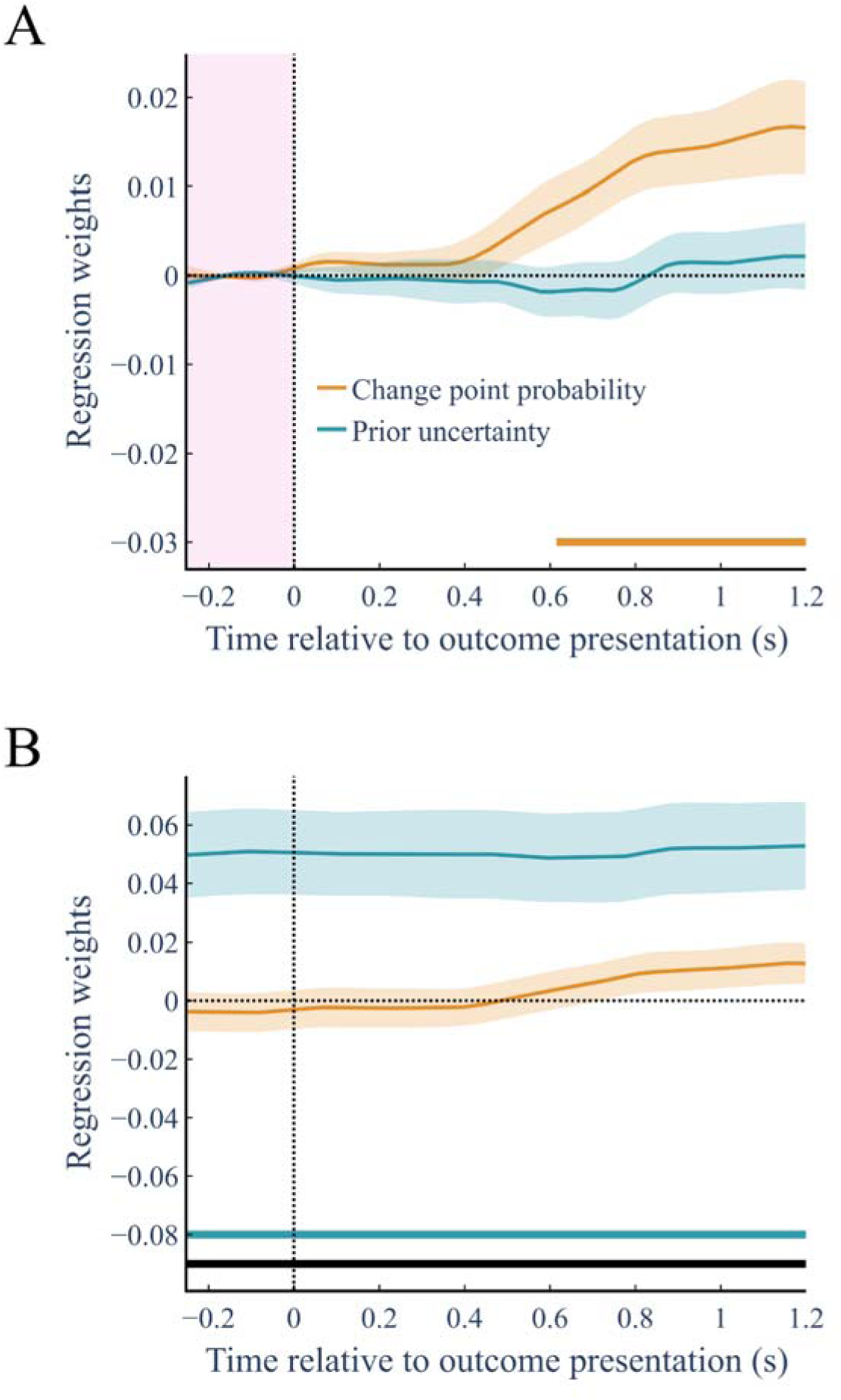
Phasic and tonic pupil diameter dynamics reflect distinct uncertainty factors within an auditory probability learning task. **(A and B)** Peristimulus regression analysis was performed on both phasic (**A)** and tonic **(B)** pupil diameter responses, as defined in the main text. The baseline for the phasic response (-250 to 0 ms) is represented on the plot (**A**). We demonstrate that the coupling between pupil signals and uncertainty factors effects holds in an auditory probability learning task of similar structure. Horizontal lines display significant time points for the corresponding (color-coded) uncertainty parameter based on cluster-based permutation analysis (*p* < 0.05, two-tailed, FWE cluster-corrected for multiple comparisons across time). Black horizontal line represents significant time points for a difference between uncertainty parameters based on cluster-based permutation analysis (*p* < 0.05, two-tailed, FWE cluster-corrected for multiple comparisons across time). All plots display the mean ± s.e.m.

**Fig S5.**
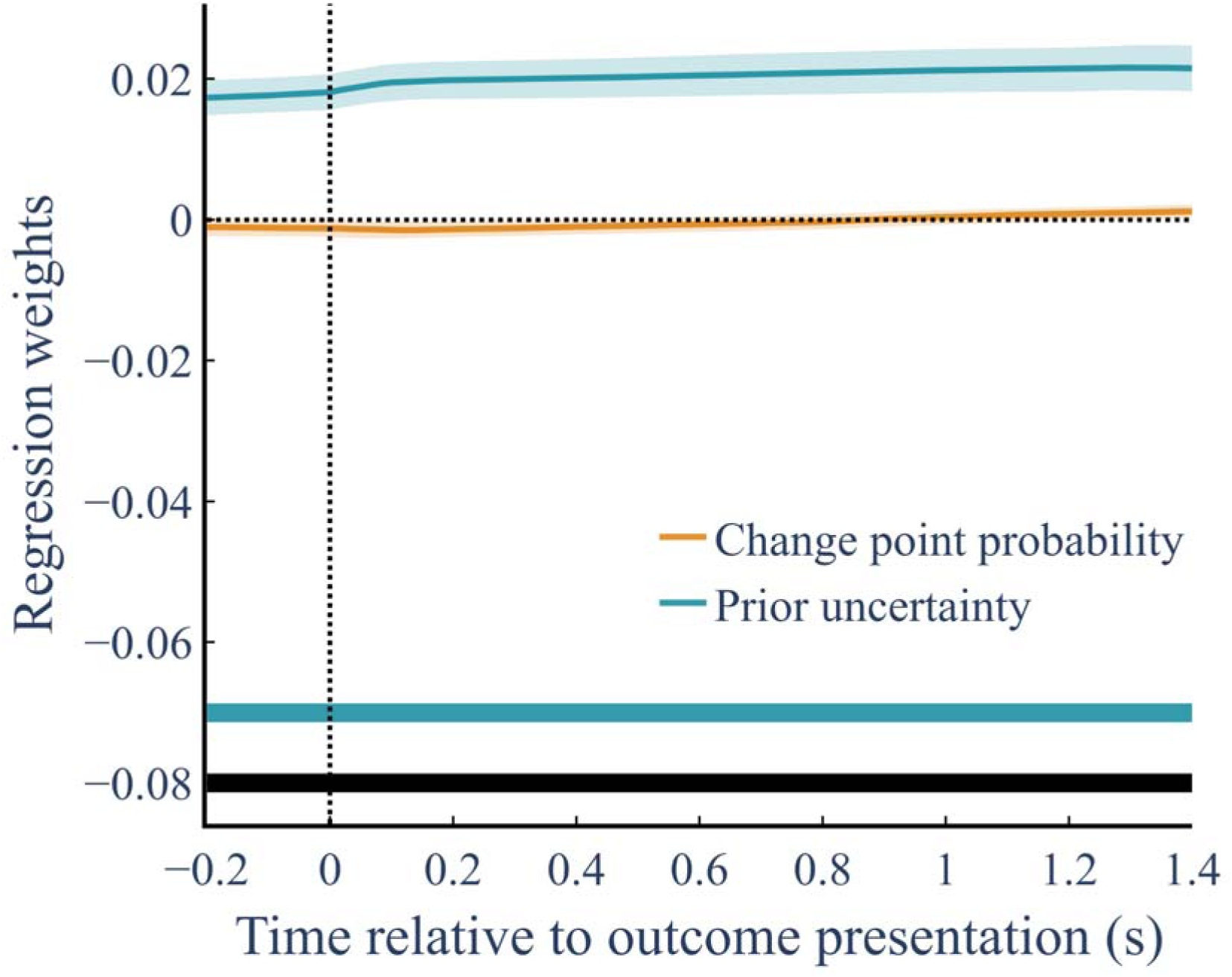
Peristimulus regression analysis performed on tonic pupil diameter derived using the deconvolution method proposed by Mittner & Groot [95]. The method adopts a Bayesian optimization approach, drawing a smooth curve (a B-spline) close to and just below prominent low points in raw pupil signals. This creates a “lower envelope” that represents the slow-moving baseline. Tonic-associated pupil responses displayed an effect of prior uncertainty across the entirety of the studied epoch. Horizontal lines display significant time points for the corresponding (color-coded) uncertainty parameter based on cluster-based permutation analysis (*p* < 0.05, two-tailed, FWE cluster-corrected for multiple comparisons across time). Black horizontal line represents significant time points for a difference between uncertainty parameters based on cluster-based permutation analysis (*p* < 0.05, two-tailed, FWE cluster-corrected for multiple comparisons across time). Plot displays the mean ± s.e.m.

**Table S1.** Slope of the linear regression between participants’ versus model estimates and learning rates.

| Parameter | Mean | s.e.m | Standard deviation | T-test against 0 |  | T-test against 1 |  |
| --- | --- | --- | --- | --- | --- | --- | --- |
|  |  |  |  | *t* | *p* | *t* | *p* |
| Probability Estimate | 1.04 | 0.02 | 0.12 | 25.96 | <0.0001 | 1.09 | 0.285 |
| Learning rate | 0.31 | 0.004 | 0.03 | 4.48 | 0.0001 | N/A | N/A |

**Table S2.** Mean and standard deviations for change point probability, prior uncertainty and average phasic pupil signal across tonic pupil signal quartiles.

| Variable | Quartile | Mean ± SD |
| --- | --- | --- |
| Change point probability | Q1 | $0.0503 \pm 0.002$ |
| | Q2 | $0.0503 \pm 0.002$ |
| | Q3 | $0.0504 \pm 0.001$ |
| | Q4 | $0.0502 \pm 0.001$ |
| Prior uncertainty | Q1 | $0.163 \pm 0.003$ |
| | Q2 | $0.168 \pm 0.002$ |
| | Q3 | $0.170 \pm 0.002$ |
| | Q4 | $0.178 \pm 0.004$ |
| Phasic pupil signal (z-score) | Q1 | $0.480 \pm 0.230$ |
| | Q2 | $0.087 \pm 0.153$ |
| | Q3 | $-0.103 \pm 0.133$ |
| | Q4 | $-0.361 \pm 0.129$ |

